# Early Progression Cells mark a stem-like state associated with pancreatic cancer initiation and progression

**DOI:** 10.64898/2026.11.03.763932

**Authors:** Beatriz Salvador-Barbero, Ella K Reed, Daniel Ruiz-Palacios, Reuben Asher, Sergio Pedraza-Arevalo, Alejandro Ibáñez-Costa, Pilar Acedo, Justo P Castaño, Catherine Hogan

## Abstract

Pancreatic cancer is deemed to progress from microscopic pancreatic intraepithelial neoplasia (PanINs). However, low grade PanINs are frequently detected in the general population without presentation or increased risk of developing cancer, suggesting that many PanINs never progress to tumours. What precisely characterises PanINs that develop into tumours is insufficiently understood. Using a mouse model where pancreatitis-induced tissue injury promotes pancreatic cancer from rare KrasG12D cells, we identify Early Progression Cells (EPCs) as a new entity, a novel population of cancer stem-like cells. Transcriptomic analyses illustrate EPCs as stress-adapted, inflammation responsive stem cells that also upregulate gene programmes indicative of microenvironment remodelling. Using identified EPC protein markers, we show that EPCs are prevalent in early metaplastic stages and in PanIN progression from low grade to high grade but are rarer in tumours. PanIN- and tumour-derived sphere and organoid assays demonstrate that EPCs possess all characteristics of stem cells in vitro and are dependent on Nupr1 stress response pathways for survival and growth. We identify APP-CD74 as a novel cell-cell interaction pathway between EPCs and neighbouring epithelial cells in PanINs and tumours. Inhibition of CD74 activity in vitro induces high-grade PanIN organoid death, suggesting EPCs are critical for high-grade PanINs. Using spatial transcriptomics of human samples, we detect gene signatures of EPCs in distinct regions of human PanIN and PDAC tissues, including APP-CD74 interactions, providing compelling evidence that EPCs are relevant to human disease. Further understanding of EPC biology will likely uncover new knowledge on why rare PanINs progress towards tumours.

## Introduction

Pancreatic ductal adenocarcinoma (PDAC) is among the deadliest of human cancers with poor survival rates (5-year survival is 10% worldwide^1^). This is mainly due to late diagnosis; although genetic studies suggest that more than ten years are needed for a genetically mutated ‘cancer-initiated’ cell to progress to metastatic PDAC^2^. Thus, there is a large window of opportunity for effective early detection and early intervention strategies, which would greatly benefit patients. Understanding the biology underlying how genetically mutant cells progress to a cancerous state is a critical first step to improving patient survival rates.

Pancreatic cancer arises sporadically from exocrine epithelial cells carrying activating mutations in the *KRAS* oncogene, which is mutated in more than 90% of human tumours^3^. Mouse studies demonstrate that the earliest metaplastic event occurs when acinar cells harbouring Kras mutations are locked in an irreversible ductal-like transdifferentiated state. In the absence of genetic mutations, Acinar-Ductal Metaplasia (ADM) is an important regenerative response of acinar cells to injury^4^. In cancer, ADM lesions harbouring mutant cells are deemed to progress towards the best-characterized precancerous lesions, the Pancreatic Intraepithelial Neoplasia (PanIN)^5,6^. Progression from a minimally dysplastic epithelium (Low Grade-PanIN; LG-PanIN) to more severe dysplasia (High Grade-PanIN; HG-PanIN) and eventually to invasive carcinoma is associated with acquisition of additional genetic mutations/aberrations^7^ and a pro-tumorigenic remodelling of the surrounding microenvironment^8^.

To recapitulate early disease stages in humans, we model sporadic pancreatic cancer in the mouse by expressing activating oncogenic KrasG12D mutations in low numbers of cells in an otherwise healthy tissue^9,10^. Using this approach, we have shown that KrasG12D cells compete for survival and are often eliminated from exocrine and endocrine tissue compartments as ‘losers’. In our model, LG-PanINs appear infrequently^9^, suggesting cell elimination processes prevent early disease development by reducing the number of oncogenic niches in tissues. Recently, we reported that some KrasG12D mutant cells override tissue protective mechanisms and persist in tissues as a heterogenous population of dormant and/or stem-like cells^10^. Persister KrasG12D cells upregulate E-cadherin-based cell-cell adhesions at mutant-normal boundaries, driven by non-canonical WNT5A signalling^10^. Remarkably, our data indicate that persister KrasG12D cells display gene signatures of pancreatic cancer even at these very early stages^10^, suggesting this population represents cancer cells of origin.

Cancer stem cells are defined as a unique subpopulation of tumour cells that possess self-renewal capabilities, chemoresistance and metastatic potential^11^. Even though cell-specific markers of Pancreatic Cancer Stem Cells (PCSCs) have been described^11–13^, their contribution to PanINs and PanIN progression are unclear. It is generally accepted that KRAS mutations alone are insufficient to drive cancer^5^; a plethora of studies show that chronic inflammation (experimental pancreatitis, as a result of tissue injury) is a potent tumour promoting factor when combined with tissues mutant for *KRAS*^5,12,13,19,20^. Based on our previous results, we hypothesised that, in the context of sporadic disease where normal-mutant cell-cell interactions are relevant, chronic inflammation would select for stem-like KrasG12D cells that progress towards tumours via PanIN stages.

Here, we identified a distinct subset of cells that express pancreatic stem/PCSC signatures, which we term ‘Early Progression Cells’ (EPCs). EPCs express stress-adapted, inflammation-responsive, ECM-interacting gene signatures when compared with other exocrine cells. Pseudotime analysis of single cell RNA sequencing (scRNAseq) data reveals EPCs as a potential progenitor-like population for cells in ADM, PanIN and tumours. These cells have higher stemness potential compared to non-EPC PanIN cells and express stress adapted responses. Analyses of cell-cell interaction networks between EPCs and neighbouring cells infer that APP-CD74 receptor-ligand interactions promote proliferation/cell survival pathways cells in PanINs and tumours. Using spatial transcriptomics, we detect gene signatures of EPCs in distinct regions of human PanIN and PDAC tissues, providing compelling evidence that EPCs are relevant to human disease.

## Results

### Identification of Early Disease Progression Cells (EPCs), a distinct cancer stem-like population of epithelial cells in pancreatic cancer initiation

We have previously shown that administration of low dose tamoxifen (1ug/40g body weight) to 6-8-week-old adult KC mice (Pdx1-CreERT; LSL-KrasG12D/+; Rosa26 LSL-tdRFP) induces expression of KRasG12D and Red Fluorescence Protein (RFP) reporter in 20-25% of the pancreatic epithelium^9^ (7 days post tamoxifen induction - p.i.). At 35 days post tamoxifen induction the percentage of RFP significantly decreases to ∼10% in KC tissues (KrasG12D)^9,10^. To induce the development of metaplasia, PanINs and PDAC tumours, we treated 35 days p.i. KC animals with the cholecystokinin receptor antagonist, caerulein, which causes experimental pancreatitis with widespread tissue injury and chronic inflammation^20,21^. Mice were harvested at defined timepoints representative of disease progression: ADM, LG-PanINs, HG-PanINs and tumours (Figure 1A & S1A). We also harvested tissues from untreated KC animals: 7 days p.i. represents a baseline timepoint, when KrasG12D mutations are induced stochastically in ∼20% of the tissue; at 35 days p.i., cell competition has occurred and only ∼50% of KrasG12D cells persist in the tissue^9,10^.

**Figure 1.**
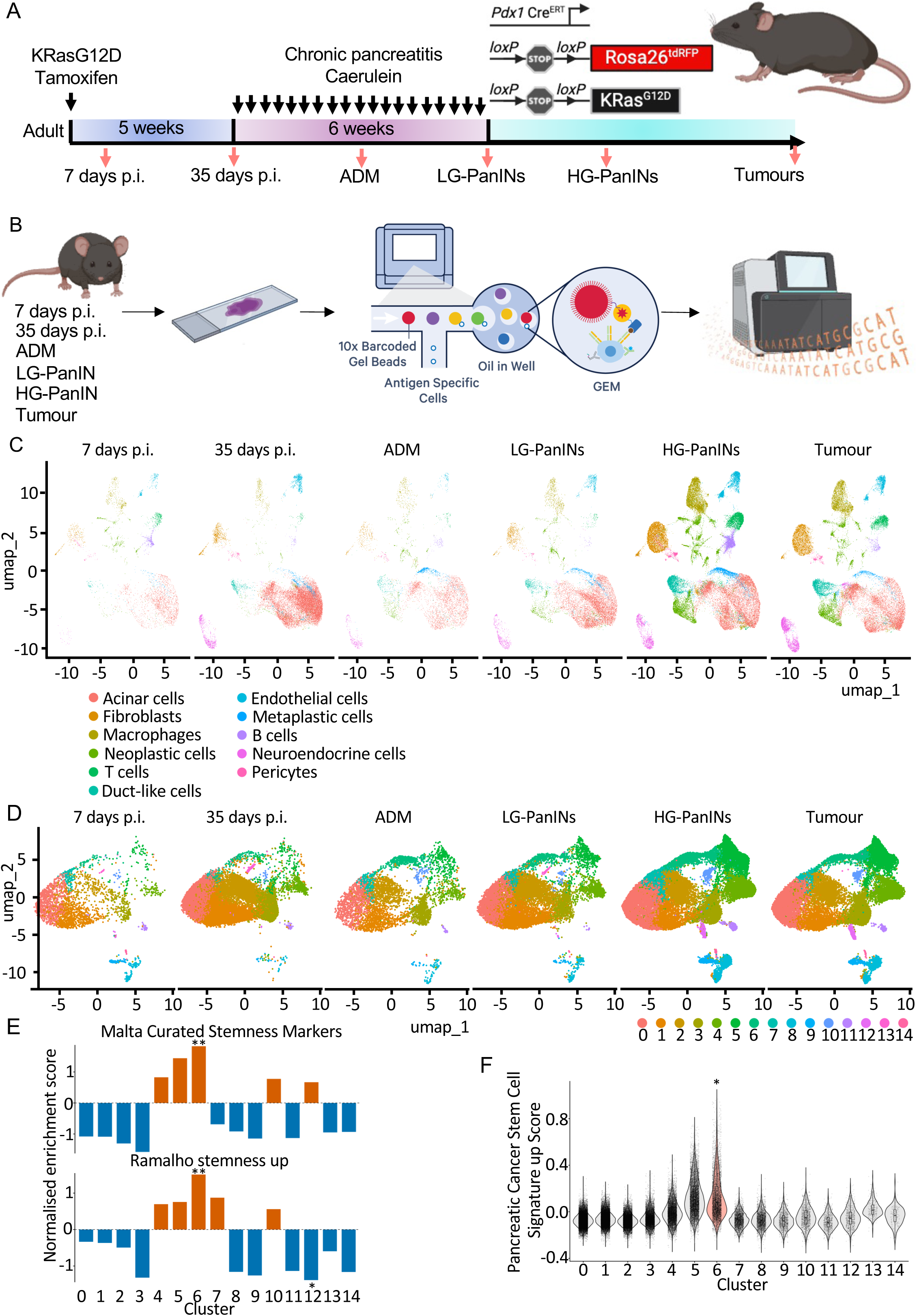
Identification of Early Progression Cells (EPCs) in single cell/nuclei transcriptomic data. (A) Schematic representation made with Biorender of the mouse model, treatment regime and timepoints. (B) Schematic representation of the scRNAseq experiment using pancreatic tissue FFPE sections. (C) Uniform Manifold Approximation and Projection for Dimension Reduction (UMAP) visualization of pancreatic scRNAseq data separated by timepoint (7 days p.i. n=2 mice; 35 days p.i. n=2; ADM n=3; LG-PanINs n=3; HG-PanINs n=3; tumours n=3) coloured according to cell annotation. (D) UMAP visualization of pancreatic exocrine cells including Acinar, Neoplastic, Duct-like and Metaplastic cells in C divided in unsupervised clusters. (E, F) Gene Set Enrichment Analysis (GSEA) of Malta_curated_stemness_markers (M30411) and Ramalho_stemness_up (MM1017) (E) and Pancreatic Cancer Stem Cell Signature Up^67^ (F) in exocrine clusters at all timepoints. Data represent mean. Wilcoxon tests were used to analyse the data. ∗P < 0.05.

We previously proposed that persister KrasG12D cells represent cancer cells of origin^9^; however, this population is heterogenous in cell fate, raising the question as to whether all remaining KrasG12D cells have potential to drive cancer development. To address this question and study disease progression, we collected tissues every 3 weeks from the start of the chronic pancreatitis regime (i.e., 8, 11, 14 and 17 weeks p.i.). To assess PanIN prevalence we performed alcian blue staining (to identify mucin producing cells^21^) and quantified number of positive lesions per area (Figure S1A, B). At 8 weeks p.i., PanINs were absent from the tissue (Figure S1A; ADM marked with dashed lines), whereas at 11 weeks p.i., PanINs were more frequent and further increased at 14 and 17 weeks p.i. (Figure S1B). To quantify the proportion of LG-PanINs versus HG-PanINs, we co-stained tissues with Mucin 5AC (present in LG-PanINs and HG-PanINs) and Mucin 1 (only present in HG-PanINs; Figure S1^23^. PanIN quantification showed a prevalence of LG-PanINs at 11 weeks p.i. and HG-PanINs at 14 and 17 weeks p.i. (Figure S1A, C). Based on these results, we established 8 weeks p.i. as ADM, 11 weeks p.i. as LG-PanINs and 14 weeks p.i. as HG-PanINs timepoints. We collected tumour tissue at humane endpoints when animals presented with tumour clinical signs. We found that endogenous RFP expression was detected in ADM, LG-PanINs, HG-PanINs and tumours tissues and the level of RFP positive area did not significantly change across timepoints (Figure S1A, D), indicating that in our model, cancer progresses from KrasG12D (RFP positive) cells.

To understand KrasG12D cell heterogeneity and determine whether a distinct population of stem-like cells are required for disease progression, we performed single-cell/nuclei RNA sequencing (Figure 1B) using FLEX GEM-X 10X Genomics technology collecting samples at each timepoint: 7 days p.i. (n=2 mice), 35 days p.i. (n=2), ADM (n=3), LG-PanINs (n=3), HG-PanINs (n=3) and tumours (n=3). Using Uniform Manifold Approximation and Projection plots (UMAPs), unsupervised clustering and population-specific gene expression for pancreatic samples previously published^12^, we annotated the different cell lineages present in our samples (Figure 1C). Briefly, we used *Resp18* to identify neuroendocrine cells, *Rgs5* for pericytes, *Ptrcp* for immune cells, *Col1a1* for fibroblasts and *Pecam1* to identify endothelial cells (Figure S2A). Within the exocrine compartment we distinguished between acinar (*Cpa1*), ductal (*Krt19*), metaplastic (*Krt19*, *Clu*) and neoplastic (*Arg2*, *Muc5ac*, *Tff1*) cells (Figure S2B). To identify the population of cells with stem-like characteristics that is associated with pancreatic cancer progression, which we call Early Progression Cells (EPCs), we focused on the exocrine compartment since cancer originates from reprogrammed acinar/ductal epithelial cells^24^, grouping together acinar, ductal-like, metaplastic and neoplastic cells for further analysis and subdivided these populations into 15 clusters (Figure 1D). We identified clusters 0, 1, 2, 3 and 7 as acinar cells, while the remaining clusters represented metaplastic/neoplastic cell states (Figure S2B). We found that previously published gastric-like cell signatures ^11,12^ were embedded in cluster 5, and cluster 10 expressed Tuft cell markers (Figure S2C).

Our previous work showed that a subset of KrasG12D cells that survive tissue protective cell elimination mechanisms, express dormant-like and stem-like characteristics^9^. We hypothesised that EPCs arise from this stem-like subset of cells, should display PCSC characteristics and be present at multiple stages of disease. To test this hypothesis and identify EPCs we used stemness, pancreatic progenitor, and PCSC signatures to cross reference with each of the clusters identified in our dataset. We found that clusters 5 and 6 expressed the highest levels of Malta_curated_stemness_markers (M30411) transcriptional signature (significantly upregulated for cluster 6 – p<0.01) with cluster 6 also significantly upregulating Ramalho_stemness_up (M9473) transcriptional signature (p<0.01 - Figure 1E). Clusters 5 and 6 showed a general increased expression of pancreatic stem/progenitor cell markers (Figure S3A) and PCSCs Signature Up^24^ (Figure 1F, S3B). We validated pancreatic progenitor/stem genes Cytokeratin 19 and CD44 were expressed at higher levels at the protein level in low numbers of cells at the different stages of disease progression (Figure S1A and S3C). We hypothesised that EPCs are crucial for early disease progression rather than tumour maintenance; therefore, we expected EPCs to exist as small populations in early precancerous lesions and persist as a minor population in established tumours. We found that cluster 5 was formed by very few cells at earlier timepoints and the number of cells dramatically increased in the later disease stages (HG-PanINs and tumours; Figure 1D & S3D). In addition, cluster 5 expressed markers of neoplastic cells present in pancreatic cancers (Figure S2C). In contrast, cluster 6 appeared as a bridge population between acinar cells (clusters 0, 1, 2, 3 and 7) and neoplastic cells (Figure 1D, S2C) and, although cluster 6 is rare in non-disease timepoints (7 and 35 days p.i.), the proportion of these cells dramatically increased at ADM timepoint, consistent with increased cell reprogramming and plasticity. At LG-PanINs, HG-PanINs and tumour timepoints cluster 6 forms a relatively small proportion of the exocrine compartment (Figure 1D & S3D), fitting with EPC dynamics. In addition, cluster 6 stemness signatures, compared to other exocrine clusters, were increased at all timepoints and along disease progression (Figure S3E)

Previous research has highlighted a high epithelial cell heterogeneity in PanINs that could be relevant for the development of pancreatic cancer initiation^11,12,25^. Within this cell heterogeneity, are subsets of cells with higher potential to drive malignancy. However, these experiments were performed using mouse models where Kras mutations are expressed in development and in most of the exocrine compartment. To understand if our data recapitulate previous findings, or identify new stem-like signatures, we compared signatures in clusters 5 and 6 to previously reported signatures. Reyes *et al* found a progenitor-like population with a defined cell state at the transition between PanINs and tumours^13^ (expression signature is shown in Figure S4A). When studying this signature in our data, cluster 6 expressed the highest levels of progenitor-like marker *Cldn4* in a relevant percentage of cells (>40%). High expression levels of *Itga3*, *Itgb4* and *Mdm2* were shared between clusters 5 and 6. However, all other markers that defined Reyes *et al* progenitor-like cells were expressed either with low average expression or in a small percentage of clusters 5 and 6 cells (Figure S4A). Similarly, genes expressed by incipient malignant cells’ MP10 programme found to drive pancreatic cancer initiation by Zhuo *et al*^21^ were expressed in low levels or low numbers of cells in clusters 5 and 6, except for *Itgb4* which is highly expressed in both clusters (Figure S4B). Our results suggest that some stem-like cells (including EPCs) identified in our model share gene expression patterns with progenitor-like/MP10 programme cells; however, the overlap between the two populations is limited.

In summary, by modelling pancreatitis-induced pancreatic cancer from rare KrasG12D cells in the context of normal cells, we found two new populations of stem-like cells, clusters 5 and 6, that are distinct in gene signatures compared to previously published data. Since cluster 6 showed increased stemness signatures and population dynamics concordant with cells relevant for disease progression rather than tumour maintenance, we propose that cluster 6 represents a ‘cancer driver/progression cell’ or EPC. With this in mind, we set out to further characterise the biology and understand the regulation of EPCs.

### EPCs expression patterns show stress-adapted, inflammation-responsive, ECM-interacting signatures with strong adhesion and survival signalling

To characterise EPCs, we studied the most Differentially Expressed Genes (DEGs – padj<0.0001) between cluster 6 (EPCs) and all the other exocrine cells. EPCs expressed genes involved in the activation of a stressed/activated epithelial programme, showing a strong tissue injury and epithelial repair signature (*Cldn4, Krt7, Krt8, EphA2, Itgβ6, LamA3, LamC2, LamB3, Thbs1*)^22,23^, extracellular matrix remodelling (*Thbs1, LamA3, LamC2, LamB3, F3, Ccn1*), cellular stress and MAPK signalling (*Dusp10, Dusp4, Ier3, Gadd45A, Cdkn1a*)^31,32^, TNF/IL1/inflammatory signalling (*Tnf, Il1rn, Tnfrsf12a, Tnfrsf23, Tnip3, Lif*), TGF-β activation and fibrosis-related signalling (*Itgβ6, Thbs1, Angptl4, F3*) and hypoxia and wound healing signatures (*Angptl4, Gdf15, Sca-1, Lif*)^37,38^ (Figure 2A and Supplementary table 1). In addition, the percentage of cells and the expression levels of these markers, increased with disease progression in cluster 6 and no other clusters (Figure S5A). To validate the presence of these markers in tissues we co-stained KC tissues harvested at ADM, LG-PanINs, HG-PanINs and tumour timepoints with antibodies against Integrin β6 and Claudin 4 (Figure 2B), since these genes were found to be among the top upregulated genes in EPCs compared to other exocrine cells (Figure 2A and Supplementary table 1). At the protein level, cells undergoing ADM seem to express these markers (Figure 2A), consistent with an initial expansion of EPCs following tissue injury (cluster 6; Figure S3D). Quantification of cells expressing high levels of Integrin β6 and Claudin 4 (Figure 2B; white arrows) per PanIN or gland-like structure (GLS) in tumours, showed that EPCs were a rare population of epithelial cells in PanINs and tumours (less than 8% in LG-PanINs, 10% in HG-PanINs and 5% in tumours; Figure S5B-C). EPCs are absent or infrequent (<10% of cells per PanIN/GLS) in at least 70% of lesions (Figure 2C-D). Less than 20% of LG-PanINs and HG-PanINs and none of the GLS in tumours were composed of EPCs above 30% (Figure 2C-D). Together, these data show that while EPCs in ADM are quite widespread, only a small proportion of PanINs harbour high numbers of cancer-progression/driver cells (EPCs) whereas these cells are rarely found in tumours, suggesting not all EPCs progress towards PanINs and tumours.

**Figure 2.**
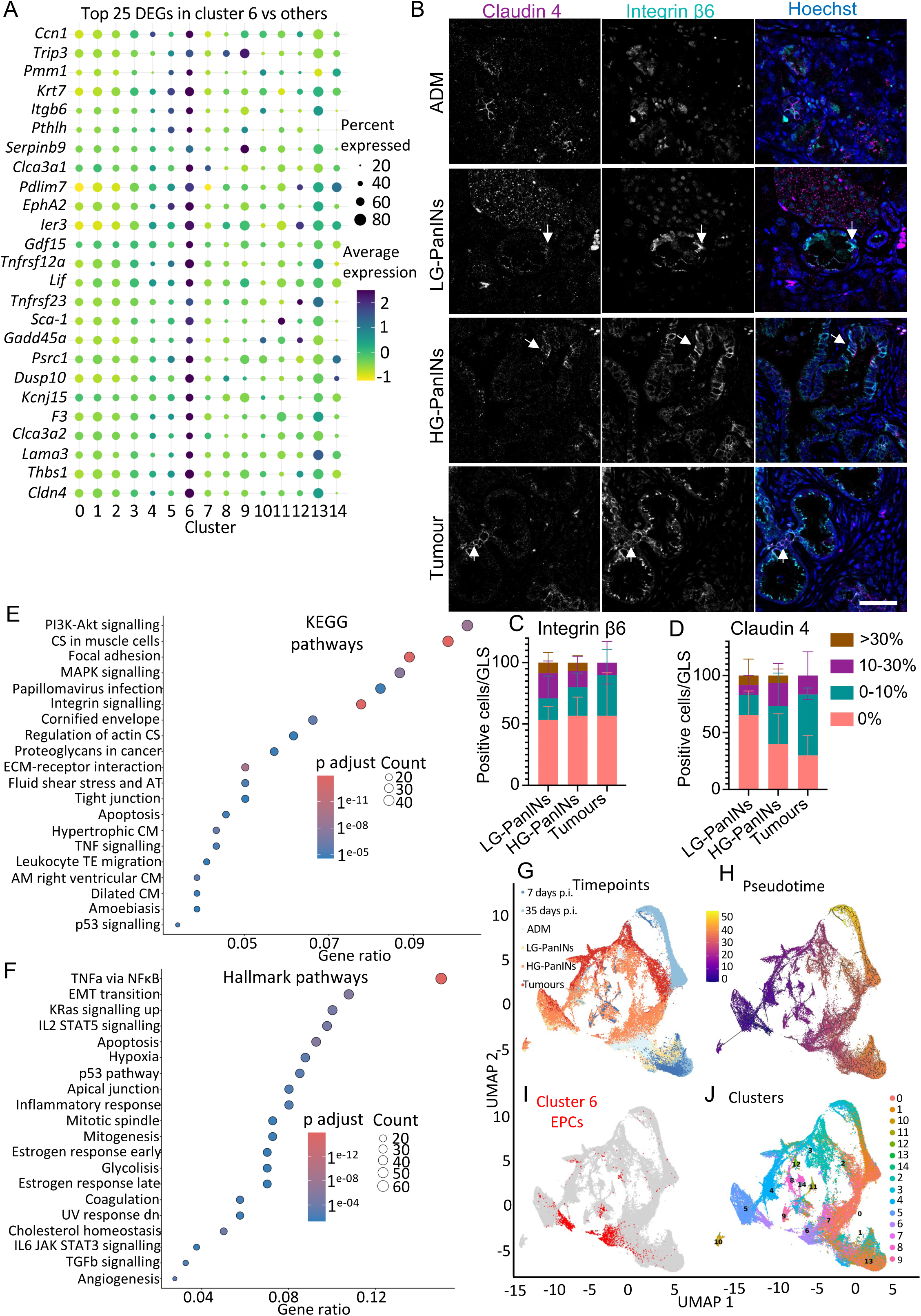
Cluster 6 (EPCs) expresses markers of cancer stem-cells and display population dynamics indicative of a stem-like population. (A) Bubble plot of the top 25 Differentially Expressed Genes (DEGs) in Cluster 6 compared to other exocrine clusters at all timepoints. Dot size represents the percent of cells in each cluster expressing the gene and the colour the log-averaged expression value. (B) Representative confocal images of mouse tissue stained with Integrin β6 and Claudin 4 at different timepoints. Scale bar: 50μm. (C-D) Bar plots showing the percentage of PanINs/GLS containing different proportions of epithelial cells showing positive staining for Integrin β6 (C) or Claudin 4 (D). Data represent mean ± SD. N=3 mice per timepoint and a minimum of 10 PanINs/GLS per mouse. (E-F) Dot plot showing significantly enriched KEGG (E) and Hallmark (F) pathways in cluster 6 compared with other exocrine clusters. Gene Ratio shows the proportion of genes associated with each pathway. Dot size indicates the number of genes contributing to each pathway (Count), while dot colour represents the adjusted p-value (Benjamini–Hochberg correction). Only pathways with an adjusted p-value < 0.0001 are shown. AT atherosclerosis; AM Arrhythmogenic; TE trans-endothelial; CM cardiomyopathy; CS cytoskeleton. (G-J) UMAP plots showing the unsupervised clustering of exocrine cells (Figure 1D), with the corresponding timepoint (G), Monocle3-inferred pseudotime trajectory, with cells coloured according to their position along the inferred trajectory from early (purple) to late (yellow) pseudotime (H), cluster 6 highlighted in red (I) and each cluster indicated by a distinct colour (J).

To characterise further the identity of EPCs, we used Gene Set Enrichment Analysis (GSEA) comparing EPCs to other exocrine cells in the transcriptomic data. We confirmed activation of pathways related to cell adhesion and ECM remodelling found in the top DEGs, e.g., focal_adhesion (M7253), integrin_signalling (NT06548) and ECM-receptor_interaction (M7098) KEGG pathways (Figure 2E), Extracellular_matrix_organisation (M610) Reactome pathway (Figure S5D), Focal_adhesion_PI3K_Akt_mTOR signalling (M39719), Focal_adhesion (M39402), Alpha6_beta4 _signalling (M39503) and Integrin_mediated_cell_adhesion (M39577) Wikipathway (Figure S5E), cell_substrate_adhesion (M15640), extracellular_structure_organisation (M18884) GO Biological process (Figure S5F). The activation of these pathways suggests EPCs are likely to be strongly anchored to and actively remodelling the local microenvironment.

We found EPCs activate inflammatory signalling pathways such as TNFa_via_NFκB (M5890), inflammatory_response (M5932) and IL6-JAK-STAT3_signalling (M5897) Hallmarks pathways (Figure 2F) and TGFb_receptor_signalling (M39351) and inflammatory_response_pathway (M39641) Wikipathways (Figure 2E and S5E).

EPCs also expressed stress-related signatures including hypoxia (M5891) Hallmark pathway (Figure 2F) and apoptosis pathways in Hallmarks, Reactome pathways, WikiPathways and GO Biological processes (M5902, M15303, M39654, M14106 and M11317; Figure 2F, S5D-F), suggesting hypoxia and environmental responses.

Pathway analysis showed that EPCs expressed gene signatures associated with pro-survival and proliferation signalling, e.g., upregulation of PI3K-AKT (mmu04151), MAPK (M10792), p53 (M6370) signalling KEGG pathways (Figure 2E), Kras_signalling_up (M5953), p53 (M5939) Hallmark pathways (Figure 2F), MAPK_family_signalling (M27565) and RAF_independent_MAPK1/3 activation (M26917) Reactome pathways (Figure S5C), EGFR1 (WP1323), MAPK (WP382), p53 (MM15852) signalling WikiPathways (Figure S5D) and ERK1_and_ERK2_cascade (M16677) GO Biological Processes (Figure S5E).

Finally, EPCs exhibited enrichment of wound healing and tissue repair-associated programmes, including wound_healing (M12074) and regulation_of_response_to_wounding (M13178) GO Biological Processes together with inflammatory response, IL6-JAK-STAT3, TGFβ and growth factor signalling pathways mentioned before (Figure 2E and S5D) and RTK_receptor_signalling (M27870) and regulation_of_IGFBPs (M27285) Reactome pathways (Figure 2E-F and S5C-E). These findings, combined with the activation of ECM remodelling and integrin-mediated adhesion pathways, suggest that EPCs adopt a coordinated tissue repair-like phenotype in addition to responding to environmental stress. The transcriptomic data also indicated that EPCs showed significantly increased Kras signalling and MAPK pathway activation (Figure 2F), supporting the interpretation that this population is enriched for KrasG12D cells.

### Trajectory and functional enrichment analysis identifies EPCs as a candidate progenitor-like population

Next, we aimed to test if EPCs behave like an early/intermediate population that could give rise to more differentiated clusters. We applied Monocle3^28,29^ trajectory inference framework to analyse exocrine populations in our scRNAseq data. We used EPCs as root point to interrogate if other clusters follow from it and test if EPCs are a progenitor/stem-like cell state. Monocle3 assigned a pseudotime value to each exocrine cell; low pseudotime values apply to cells closer to the root transcriptional state while high pseudotime values show cells further along the inferred trajectory. As expected, pre-pancreatitis timepoints showed the most differentiated states (i.e., higher pseudotime values; Figure 2G timepoints and pseudotime panels, and S6A). The ADM timepoint represented an intermediate state between pre-pancreatitis (7 and 35 days p.i.) and later timepoints (Figure 2G-H and S6A). In LG-PanINs timepoint tissues, we observed PanINs surrounded by histologically normal exocrine cells (Figure S1A; LG-PanIN panel), suggesting most acinar cells resolve to adopt a differentiated state post caeureilin exposure. The pseudotime analysis supported this observation, with most exocrine cells clustering with earlier timepoints (7 and 35 days p.i. and ADM; Figure 2G-H and S6A). In contrast, some cells in LG-PanINs timepoint positioned closer to cells in HG-PanINs and tumours timepoints with lower pseudotime values (Figure 2G-H and S6A), suggesting some PanIN cells transdifferentiate towards a metaplastic state at this timepoint. At HG-PanINs and tumour timepoints, we observed a wide range of pseudotime values (Figure 2G-H and S6A), suggesting high heterogeneity in differentiation states between cells present in tissues at these timepoints.

Cluster 5 was predominantly associated with low pseudotime values, whereas cells within cluster 6 (EPCs) spanned a broad range of pseudotime values despite this population containing the trajectory root (Figure 2H-J and S6B). This suggests that EPCs are divided into (at least) 2 major subpopulations: one with the lowest pseudotime values that branched initially into cluster 5, then 4 and later into more differentiated clusters such as 3, 12 and 8 (corresponding with a more neoplastic population containing cells from later timepoints; HG-PanINs and tumours); and a second subpopulation, closely associated with ADM timepoint, that branched out into clusters 7, 0, 1 and 2; the acinar cell clusters. Our analysis infers that EPCs expand when acinar cells undergo ADM; a proportion of these cells re-differentiate to an acinar cell fate, however some retain a progenitor-like state that can, potentially, further de-differentiate to drive pancreatic cancer initiation. This suggests that EPCs are not a uniform cell cluster/population but instead, are composed of different cells at different stages along a dynamic transcriptional process, potentially reflecting its progressive diversification during pancreatic cancer progression. Gene module activity per cluster (the group of genes that show similar expression patterns across the trajectory) showed module 1 to be most upregulated in EPCs and to a lesser extent also in cluster 5 (Figure S6C). GO Biological processes analysis confirmed module 1 genes to be involved in cell dedifferentiation (epithelial_cell_development (MM4161), skin_development (M15889), lung_development (M24167); Figure S6D), supporting our conclusion that EPCs represent a progenitor/stem-like population that can give rise to different cell states.

In summary, pseudotime analysis positions EPCs as a candidate progenitor/stem-like population with a trajectory, which branches from EPCs towards acinar cell differentiation or neoplastic states, and enrichment of transcriptional modules associated with progenitor activity.

### EPCs have a higher stemness potential than other PanIN cells

To explore the stemness potential of EPCs, we first studied the top DEGs in EPCs compared to other exocrine cells (Figure 2A). The tenth most DEGs was Ly6a or Stem cell antigen 1 (Sca-1; Figure 2A) gene. Sca-1 is a cell-surface glycosylphosphatidylinositol-anchored protein associated with cellular plasticity, self-renewal capacity, tissue regeneration and resistance to stress^30^. In pancreatic cancer, Sca-1 is a marker for tumour-propagating and progenitor-like populations in mouse models^16,17,42^. Similar to other markers found to be upregulated in EPCs, Sca-1 was expressed in most cells undergoing ADM, while present only in few cells in most PanINs and GLS in tumours (percentage of total Sca-1 positive cells: 7% of cells in LG-PanINs, 13% in HG-PanINs and 5% in tumours; Figure S7B), with only a few PanINs (10% of LG-PanINs and 4% of HG-PanINs) and very rarely in GLS in tumours (3% of GLS) showing more than 30% of these cells (Figure S7A, C).

To functionally test the ability of EPCs to behave as stem-like cells, we used sphere formation assays. We first confirmed that 80% of Sca-1 positive cells in tissues were RFP positive (data not shown). Pancreas tissues of KC mice were harvested at LG-PanINs and HG-PanINs timepoints and then Mucin5AC positive cells were sorted (using Magnetic-Activated Cell Sorting; MACS) to select PanIN cells. To differentiate between EPCs and other epithelial cells in PanINs, we sorted Mucin 5AC positive cells into Sca-1 positive (EPCs) and Sca-1 negative cells. The same number of sorted cells were seeded in low attachment plates and grown for 14 days. We confirmed that all cells in the experiment were RFP positive. Sphere quantification showed a significant increase in the number of spheres formed from Sca-1 positive (EPCs) compared to Sca-1 negative cells at both LG-PanINs and HG-PanINs timepoints (p=0.0281 and p=0.0268 respectively; Figure 3A-B) and an increased sphere diameter (p=0.0054 and p=0.0005 respectively; Figure 3A, C).

**Figure 3.**
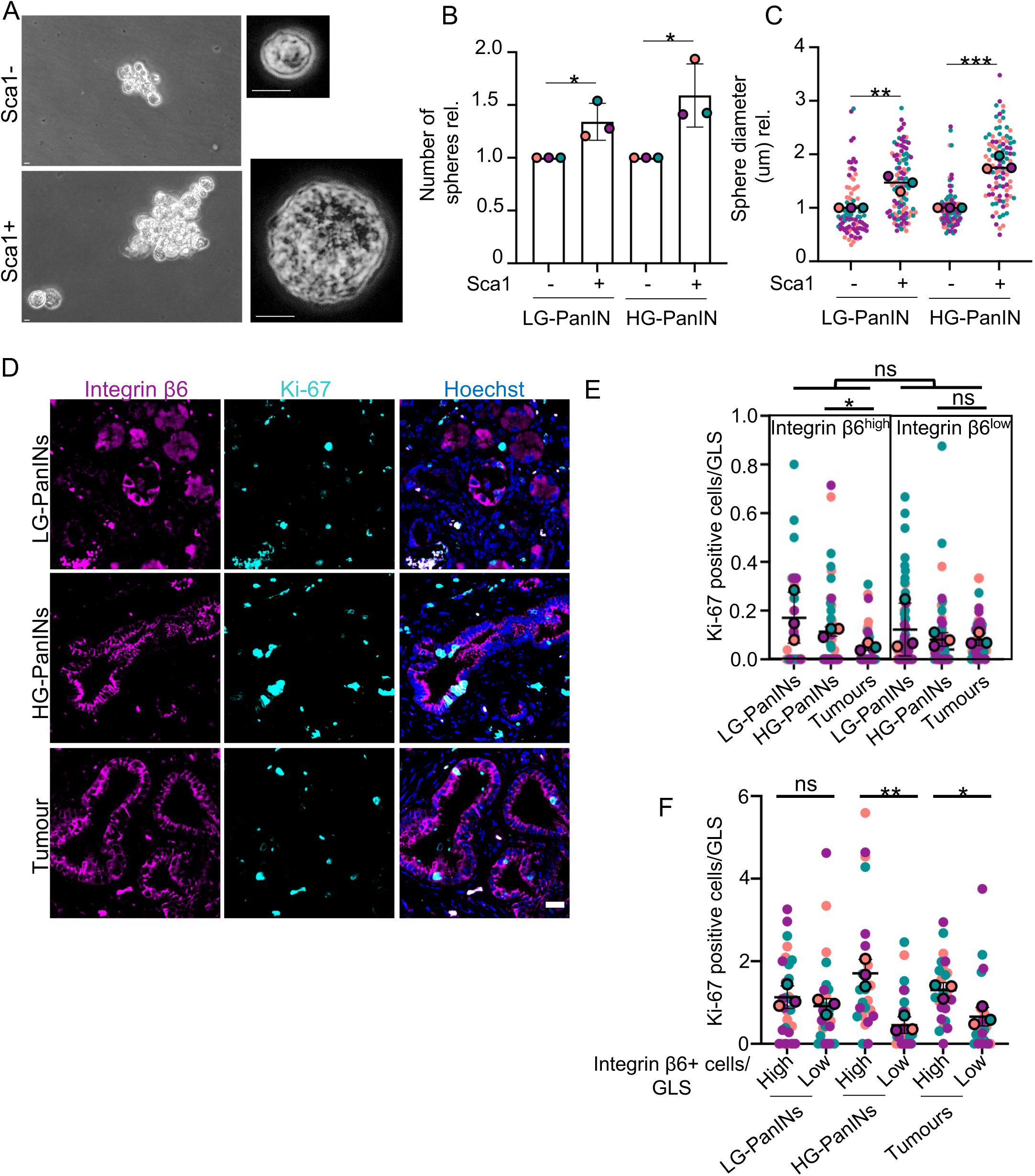
EPCs show characteristics of stem cells to a greater extent than non-EPCs. (A) Representative brightfield images of sphere formation assays using Sca-1 positive and Sca-1 negative isolated from HG-PanINs. Scale bar: 10μm. (B) Relative number of spheres generated from Sca-1 positive or Sca-1 negative cells obtained from LG-PanINs or HG-PanINs (n=3 mice). Data represent mean ± SD. (C) Graph showing relative sphere diameter from Sca-1 positive or Sca-1 negative cells obtained from LG-PanINs or HG-PanINs. Each smaller dot represents a sphere, and bigger dots represent each corresponding colour coded mouse. (D) Representative images of mouse tissue stained with Integrin β6 and Ki-67 proliferative marker at different timepoints. Scale bar: 20μm. (E, F) Graphs showing Ki-67 positive cells per PanIN/GLS at each timepoint differentiating between Integrin β6 positive and negative cells (E) and high (>30% of Integrin β6 positive cells per PanIN/GLS) or low (<30%) PanIN/GLS (F). Each smaller dot represents a PanIN/GLS, and bigger dots represent each corresponding colour coded mouse. Data represent mean. (B, C, E, F) Student t tests were used to analyse the data. *P < 0.05; **P < 0.001; ns=P > 0.05.

To understand if EPCs are proliferative and can give rise to other cells *in vivo*, we stained mouse pancreas tissues at LG-PanINs, HG-PanINs and tumour timepoints with Ki-67 (marker of cell proliferation) and Integrin β6 (to label EPCs; Figure 3D). Quantification of Integrin β6^high^ (EPCs) and Integrin β6^low^ (low or no expression) that were also positive for Ki67 showed no significant differences between EPCs and other epithelial cells in PanINs and GLS in tumours at any timepoint (p=0.6058 LG-PanINs, p=0.1804 HG-PanINs, p=0.1675 tumours; Figure 3D-E). EPC proliferation significantly decreased with disease progression from HG-PanINs to tumour timepoints (p=0.0140), while no differences were found in integrin β6^low^ cells in PanINs/GLS (p>0.9999; Figure 3D-E). We hypothesised that EPCs are drivers of early disease stages and therefore we interrogated if cell proliferation was different in PanINs/GLS containing a high (>30% EPCs: EPC^high^) or low (<30% EPCs: EPC^low^) percentage of EPCs. No differences were found in proliferation between EPC^high^ and EPC^low^ LG-PanINs (p=0.3286; Figure 3D, F). Notably, LG-PanIN timepoints coincide with the final caereulin injections (Figure1A), suggesting the high cell proliferation counts in LG-PanIN tissues may reflect tissue regeneration post injury, making data interpretation difficult. In contrast, the proliferation rate was significantly higher in EPC^high^ compared to EPC^low^ PanINs/GLS in HG-PanINs and tumours timepoints (p=0.0088 and p=0.0197 respectively; Figure 3D, F), suggesting that although EPCs are not actively proliferating, lesions containing EPCs have higher rates of cell proliferation.

In summary, we showed that EPCs have higher stemness potential than other epithelial cells in PanINs. Although EPCs are not more proliferative than other PanIN/GLS cells, HG-PanINs containing more than 30% of EPCs showed higher proliferation rates than those containing very few or no EPCs.

### EPCs show signals of stress adaptation and survival

Our transcriptomic data showed that EPCs (cluster 6) were expressing oxidative stress, hypoxia and pro-survival responses at a higher level than other epithelial exocrine cells (Figure 2E-F and S3C-E), suggesting that EPCs are more resistant to stress. To further our understanding of stress resistance of EPCs, we performed GSEA analysis of stress-related signatures in EPCs compared to other exocrine cells. Interestingly, we found that EPCs overexpressed pathways involved in stress fibre assembly, DNA damage/p53 response, oxidative and osmotic stress, intrinsic apoptosis and hypoxia (Figure 4A). We then interrogated the tissues using cellular stress response markers such as Nupr1 and Clusterin. Nupr1 is a stress sensor and mediator of cell resistance to stress in pancreatitis^34^ and has been shown to be crucial for PanIN development^35^. We found that *Nupr1* was highly expressed in acinar clusters (0, 1, 2 and 7) and neoplastic/metaplastic clusters 6 (EPCs) and 13 (Figure S2B, 4B). Tissue staining showed most exocrine cells expressed low levels of cytoplasmic Nupr1 while a minority expressed high nuclear levels (Figure 4C), suggesting a small proportion of PanIN cells (14% in both LG-PanINs and HG-PanINs; Figure S8A) activate Nupr1 stress responses, and a minority (∼ 7%) of PanINs contained high Nupr1 expression in <50% of cells (Figure 4C-D).

**Figure 4.**
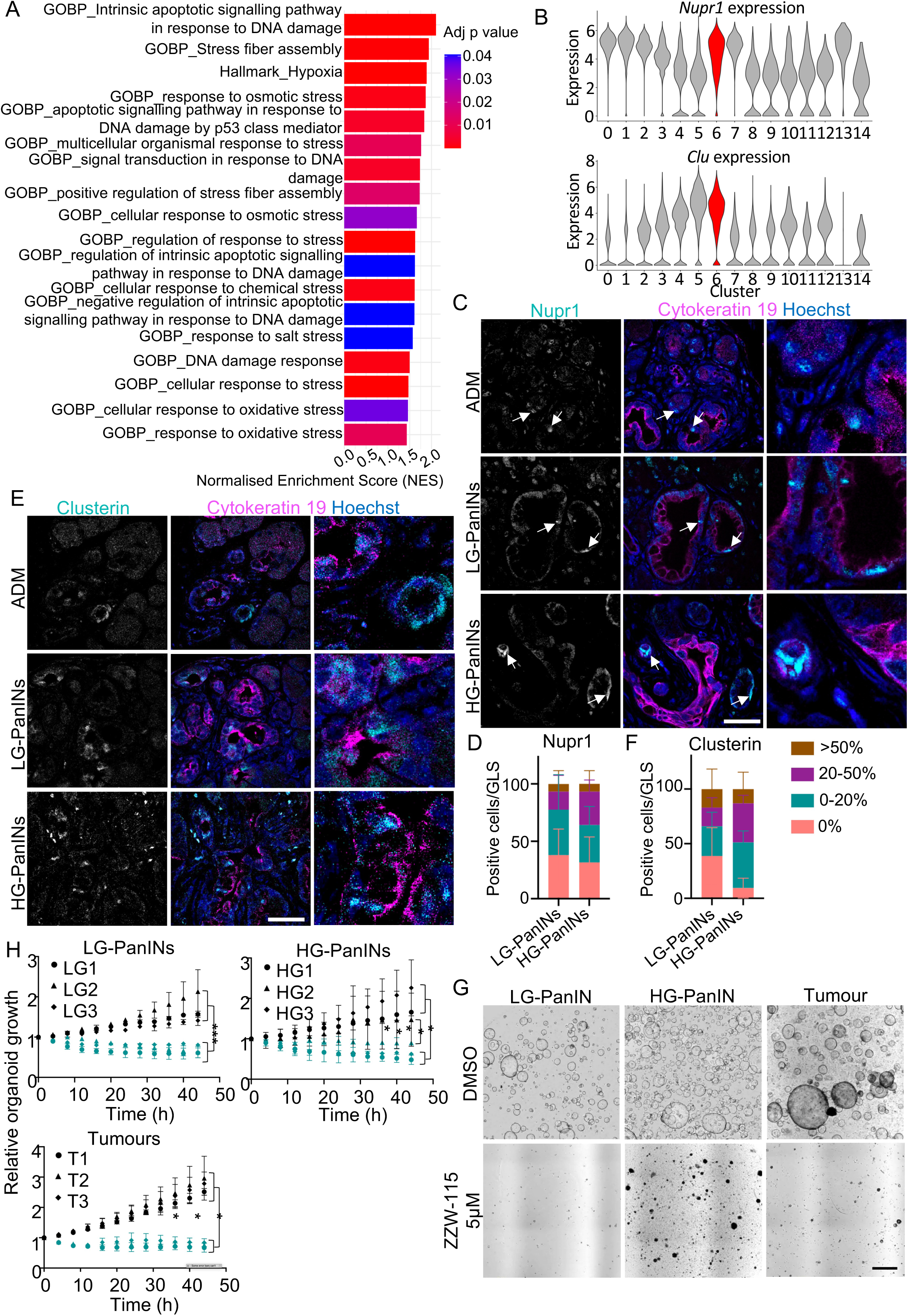
Survival of EPCs from PanINs and tumours requires stress responses. (A) Normalized Enrichment Scores (NES) of Gene Set Enrichment Analysis (GSEA) of Hallmark pathways and Gene Ontology Biological Processes (GOBP) gene sets involved in stress response in cluster 6 compared to other exocrine clusters. (B) Violin plots showing the gene expression of *Nupr1* (top) and *Clu* (bottom) stress response genes in each cluster with cluster 6 highlighted in red. (C, E) Representative confocal images of mouse tissue stained with Cytokeratin 19, Hoechst and either Nupr1 (C) or Clusterin (E) at different timepoints. Scale bar: 20μm. (D, F) Bar plots showing the percentage of PanINs/GLS containing different proportions of epithelial cells positive for Nupr1 (D) and Clusterin (F). Data represent mean ± SD. N=3 mice per timepoint and a minimum of 10 PanINs/GLS per mouse. (G) Brightfield images of LG-PanIN, HG-PanIN or Tumour organoids treated with DMSO or ZZW-115 for 48h. Scale bar: 1mm. (H) Graphs showing organoid growth over time in 3 organoid lines (3 experiments per line) obtained from 3 independent mice at each timepoint treated with DMSO or ZZW-115. Data represent mean ± SD. ANOVA tests were used to analyse the data. **P < 0.001; *** P > 0.0001.

Clusterin functions in apoptotic cell death and as a stress-induced secreted chaperone protein^36^. Clusterin is highly induced during pancreatic regeneration, particularly in ductal cells, and it is upregulated in pancreatitis increasing viability after cell stress^37^. Similarly to Nupr1, our results showed increased Clusterin gene *Clu* expression in EPCs and cluster 5 (Figure 4B). Tissue staining showed widespread staining of Clusterin in ductal-like cells, however only around one fifth of cells (19% in LG-PanINs and 24% in HG-PanINs; Figure S8B) showed high expression (Figure 4E), and only 17% of LG-PanINs and 13% of HG-PanINs contained more than 50% of cells expressing high levels of Clusterin (Figure 4E-F). Our results showed some markers of stress fibre assembly in EPCs, including Integrin β6 and Claudin 4 overexpression (Figure 2B-D). Stress-fibre assembly and actomyosin contractility enable epithelial cells to adapt their mechanical properties, maintain interactions with the extracellular matrix and respond to changes in their microenvironment. We therefore investigated whether EPCs exhibit additional molecular features that could support cytoskeletal remodelling and cellular adaptation to stress. . We next investigated EphA2, a receptor tyrosine kinase involved in cytoskeletal remodelling, cell migration and signalling pathways that can promote cell survival under adverse conditions^39^. We found *EphA2* expressed in the top 25 DEGs in EPCs (Figure 2A). Tissue staining for phospho-EphA2 (pEphA2), the active form of EphA2, showed prominent membrane staining in PanINs and not in ADM; however, only a small proportion of PanIN cells (16% in LG-PanINs and 20% in HG-PanINs; Figure S8E) in PanINs/GLS expressed high levels (Figure S8F). We found that less than 10% of PanINs contained a high proportion (>50%) of pEphA2 positive cells (Figure S8D-E).

To test whether stress responses activated in EPCs are critical for disease progression, we treated organoids derived from mouse tissues at LG-PanINs, HG-PanINs and tumour timepoints with a Nupr1 inhibitor. Reassuringly, PanIN/tumour-derived organoids were consistently positive for Nupr1 expression (Figure S8F). To test the effect of Nupr1 inhibition in PanINs and tumours we seeded three organoid lines (obtained from three independent mice) harvested at each timepoint. After 24h, organoids were treated with ZZW-115^40^ or DMSO control. Our results showed that independent of timepoint, organoid growth was impaired upon Nupr1 inhibition (LG1, LG2, LG3, HG1, HG3, T1, T2 and T3 p<0.0001 and HG2 p=0.0013; Figure 4G-H), suggesting that Nupr1 is critical for the survival of PanIN and tumour cells and for organoid growth.

In summary, EPCs upregulate cell stress-related pathways and stress response markers Nupr1 and Clusterin. Stress response inhibition with a Nupr1 inhibitor induced cell death in organoids derived from tissues representative of all disease stages. Therefore, we conclude that Nupr1 mediated cell stress responses are essential for PanIN and tumour derived organoid growth and survival.

### EPCs interact with other epithelial cells through APP-CD74 interactions. CD74 expressing cells activate inflammatory and pro-survival signature in pancreatic cancer initiation

Epithelial cell-cell interactions are essential for pancreatic cancer initiation^12,13^. We hypothesised that cell-cell interactions between EPCs and neighbouring epithelial cells are important in disease progression. To address this question, we interrogated ligand-receptor interactions in scRNAseq data using CellChat^41^. CellChat infers and analyses cell-cell communication networks by identifying ligand-receptor pairs. We investigated interactions between cluster 6 (EPCs) (as ligand expressing), and any other epithelial cell cluster (including EPCs; as receptor-expressing). Consistently, EPCs interacting with EPCs (cluster 6->6; Figure 2E-F and S5C-E) identified pathways involved in cell junction regulation [*Ocln-Ocln, F11r-F11r, Cdh1-*(*Itga1+Itgb1*)*, Cdh1-Cdh1, Lama5-*(*Itga2+Itgb1*)*, Lama5-*(*Itga6+Itgb4*)*, Lama5-*(*Itga6+Itgb1*)], tissue adhesion and cohesion [*Thbs1-*(*Itga3+Itgb1*)*, Lama5-*(*Itga3+Itgb1*)*, Spp1-*(*Itga4+Itgb1*)*, Ins-Inssr*], wound repair [*Tbsh1-Cd74, Thbs1-Sdc4, Thbs1-Sdc1, Lgals9-Havcr2, Spp1-*(*Itga4+Itgb1*)*, Spp1-Cd44, App-Cd74*], tissue integrity and cell communication (*Lama5-Cd44, Lgals9-Cd44*), and stem cell and differentiation (*Jag1-Notch1, Jag1-Notch2*) (Figure 5A).

**Figure 5.**
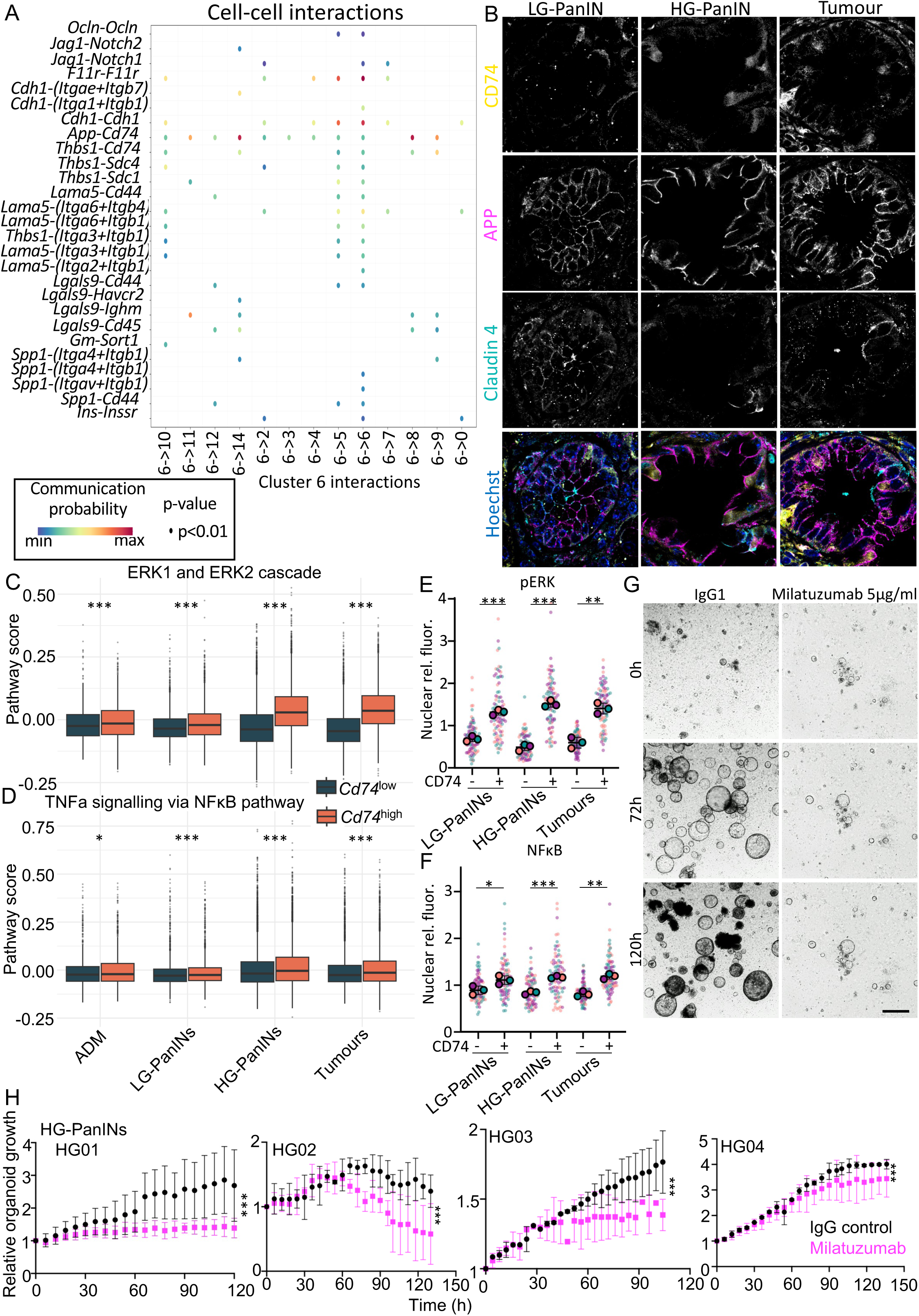
EPCs interact with other epithelial exocrine cells through APP-CD74. (A) Bubble plot showing predicted ligand-receptor interactions between Cluster 6 and the indicated receiver exocrine cell populations. Each row represents a ligand-receptor pair, and each column represents a receiver cell population. Bubble size indicates the statistical significance of the interaction, while colour represents the predicted communication probability, reflecting the strength of the inferred interaction. Only statistically significant interactions are shown. (B) Representative confoca? images of mouse tissues stained for CD74 and APP at different timepoints. Scale bar: 20μm. (C, D) Box plots showing pathway scores for the GOBP_ERK1_and_ERK2_cascade (C) and TNFa_via_NFκB_pathway (D) in *Cd74*^high^ (top quartile) and *Cd74*^low^ (bottom quartile) exocrine cells at each timepoint. Wilcoxon tests were used to analyse the data. (E, F) Graphs showing phospho-ERK (pERK; E) and NFκB (F) nuclear fluorescence intensity in CD74 positive or negative cells at each timepoint. Each smaller dot represents a PanIN/GLS, and bigger dots represent each corresponding colour coded mouse. Data represent mean values. Student t tests were used to analyse the data. (G) Brightfield images of HG-PanIN organoids treated with IgG1 or Milatuzumab 5μg/ml for 0, 72 and 120h. Scale bar: 1mm. (H) Graphs showing organoid growth over time in four independent HG-PanIN organoid lines (3 independent experiments per line) treated with IgG1 or Milatuzumab for 90-148h. Data represent mean ± SD. ANOVA tests were used to analyse the data. (C-F, H) *P > 0.05; **P < 0.001; *** P > 0.0001.

The data showed that EPCs largely interact with clusters 5, 6 and 10, except for *Lgals9* ligand interactions, which are more common with clusters 8, 9, 11, 12 and 14, and Spp1 ligand interactions with clusters 6 and 5 (Figure 5A). The most shared ligand-receptor epithelial-epithelial cell interaction between EPCs and all the clusters was Amyloid-beta precursor protein (*App*)-*Cd74* (Figure 5A). APP-CD74 has been reported to mediate endothelial cell-macrophage communication^42,43^, monocytic myeloid-derived suppressor-B cell interactions^44^, epithelial cell-macrophage crosstalk^45,46^ and epithelial cell-cancer associated fibroblasts^47^, among others. Epithelial-epithelial cell APP-CD74 crosstalk has been described between malignant cells in testicular tumours, with *APP* expressing cells having a more immunosuppressive transcriptome and suggesting that APP-CD74 interactions might drive immunosuppression^48^. A role of APP-CD74 cell-cell interactions in non-cancerous epithelial tissues is otherwise poorly described. We confirmed the expression of *App* in EPCs in our transcriptomic data and found that *App* expression increased with disease progression (Figure S9A). Immunostaining of mouse tissues at different timepoints, confirmed that APP was highly expressed in the membrane of epithelial cells in some PanINs/GLS in tumours and EPCs (as indicated by claudin 4 positivity) express APP (Figure 5B). CD74 positive cells were observed only in PanINs/GLS containing APP expressing cells (Figure 5B).

We hypothesised that APP binding activates CD74 downstream pathways in the receiving cell. In cancer, CD74 has been shown to interact with the macrophage migration inhibitory factor (MIF) and using CD44 as co-receptor, activates ERK1/ERK2 MAPK and PI3K/AKT activation^49–51^. In in B cell lymphoma, CD74 can lead to NFκB pathway activation ^49,51^. To determine if these pathways were activated in CD74 positive epithelial cells, we compared the expression of GOBP_ERK1_and_ERK2_cascade, Hallmark_TNFA_signalling_via_NFκB and Hallmark_PI3K_Akt_mTOR_signalling pathways in the top 25% (*Cd74*^high^) and bottom 25% (*Cd74*^low^) *Cd74* expressing exocrine cells at each of the different timepoints. We found ERK1/ERK2 pathway to be significantly upregulated in *Cd74*^high^ compared to *Cd74*^low^ cells in HG-PanINs and tumours timepoints (p<0.0001 at all timepoints Wilcoxon test; Figure 5C). We then looked at the positive and negative regulation of ERK1/ERK2 pathway and found GOBP_positive_regulation_of_ERK1_and_ERK2_cascade to be highly and significantly upregulated at all timepoints (median difference: ADM 0.0156; LG-PanINs 0.0246; HG-PanINs 0.1086; tumours 0.1350 and p<0.0001 at all timepoints Wilcoxon test; Figure S9B top panel). GOBP_negative_regulation_of_ERK1_and_ERK2_cascade was significantly upregulated at ADM, HG-PanINs and tumours timepoints (median difference and p value: ADM 0.0008, p=0.000978; LG-PanINs 0.0069, p=0.249; HG-PanINs 0.0258, p<0.0001; tumours 0.0089, p=0.00275 Wilcoxon test; Figure S9B bottom panel). Together, these data suggest a state of sustained ERK pathway activation coupled to transcriptional feedback regulation, consistent with dynamically controlled MAPK signalling.

NFκB analysis showed a significant increase in the activation of the pathway in Cd74^high^ compared to Cd74^low^ cells at all timepoints (ADM p=0.0294, LG-PanINs p=0.0004, HG-PanINs p<0.0001and tumours p<0.0001 Wilcoxon test; Figure 5D). Expression of PI3K/AKT pathway indicated a general but non-significant increase in Cd74^high^ compared to Cd74^low^ epithelial cells in ADM, LG-PanINs and HG-PanINs, and a significant decrease at tumour timepoint (ADM p=0.0643, LG-PanINs p=0.979, HG-PanINs p=0.515 and tumours p=0.00606 Wilcoxon test; Figure S9C), suggesting CD74 expressed on exocrine cells is not signalling via the PIK/AKT pathway.

To confirm the results at the protein level, we co-stained mouse tissues with CD74 and phospho-ERK (pERK), or CD74 and NFκB. We found that CD74 positive cells in PanINs had significantly higher expression of nuclear pERK in tissues harvested at LG-PanINs, HG-PanINs and tumours timepoints (LG-PanINs p=0.0003; HG-PanINs p<0.0001; tumours p=0.0015; Figure 5E, S9D), consistent with the transcriptomic data. In addition, we found that NFκB was significantly increased in the nuclei of CD74 positive cells compared to CD74 negative cells at all timepoints (LG-PanINs p=0.0436; HG-PanINs p=0.0004; tumours p=0.0011; Figure 5F, S9E). These data suggest CD74 positive cells activate pro-survival/anti-apoptotic signalling pathways and pro-inflammatory cell responses in PanINs and tumours.

To test the functional requirement of CD74 as a pro-survival signal in pancreatic tumorigenesis, we generated organoids from mouse tissues harvested at LG-PanINs, HG-PanINs and tumours timepoints and treated them with the anti-CD74 antibody milatuzumab^52^ or an IgG control. We consistently found that HG-PanIN-derived organoids significantly decreased in growth after milatuzumab treatment (HG01, HG02, HG03, HG04 p<0.0001; Figure 5G-H), suggesting that CD74 activation is critical for survival and growth of organoids derived from HG-PanIN tissues. In contrast, the results from LG-PanINs and tumours organoids were inconsistent (Figure S10A), with some lines showing growth inhibition while others not. These results likely reflect a high heterogeneity in cell composition and CD74 expression found in LG-PanINs/GLS in tumours. In contrast, organoids derived from HG-PanIN tissues may be more uniform for CD74 expression and hence activity.

To investigate the relevance of our observations in mouse in the human context, we performed CosMx Human Universal Characterisation Panel (HUCP) in human tissues containing primary treatment naïve tumours and normal-adjacent tissues with PanINs (n=2) annotated by a pathologist. To test the presence of EPCs, we interrogated 3 of the top 25 DEGs identified and validated at the protein level in mouse tissues (Figure 2A-B, S8F) and available in the HUCP: *CLDN4*, *EPHA2* and *ITGΒ6*. We found that in general, *CLDN4*, *EPHA2* and *ITGΒ6* gene expression was low in most cells in PanINs; however, some PanINs contained areas where a high proportion of cells expressed these genes (non-EPCs and EPCs regions respectively; Figure 6A-B top panels). EPC gene markers were expressed in single cells sparsely distributed in tumours as well as in cell clusters (Figure 6C-D top panels). To confirm the presence of EPC markers at the protein level, we stained consecutive tissue slices to the spatial transcriptomic slice with Integrin β6. Guided by the spatial transcriptomics data (Figure 6A-B), we found a consistent high level of Integrin β6 staining in PanIN regions identified as EPC-containing at the gene level, and low staining in the non-EPCs region (Figure 6E). Similarly, we found in tumours (consecutive slice to spatial transcriptomics) clusters of cells with a strong co-staining for integrin β6, as well as weakly stained cells for both makers (Figure 6E).

**Figure 6.**
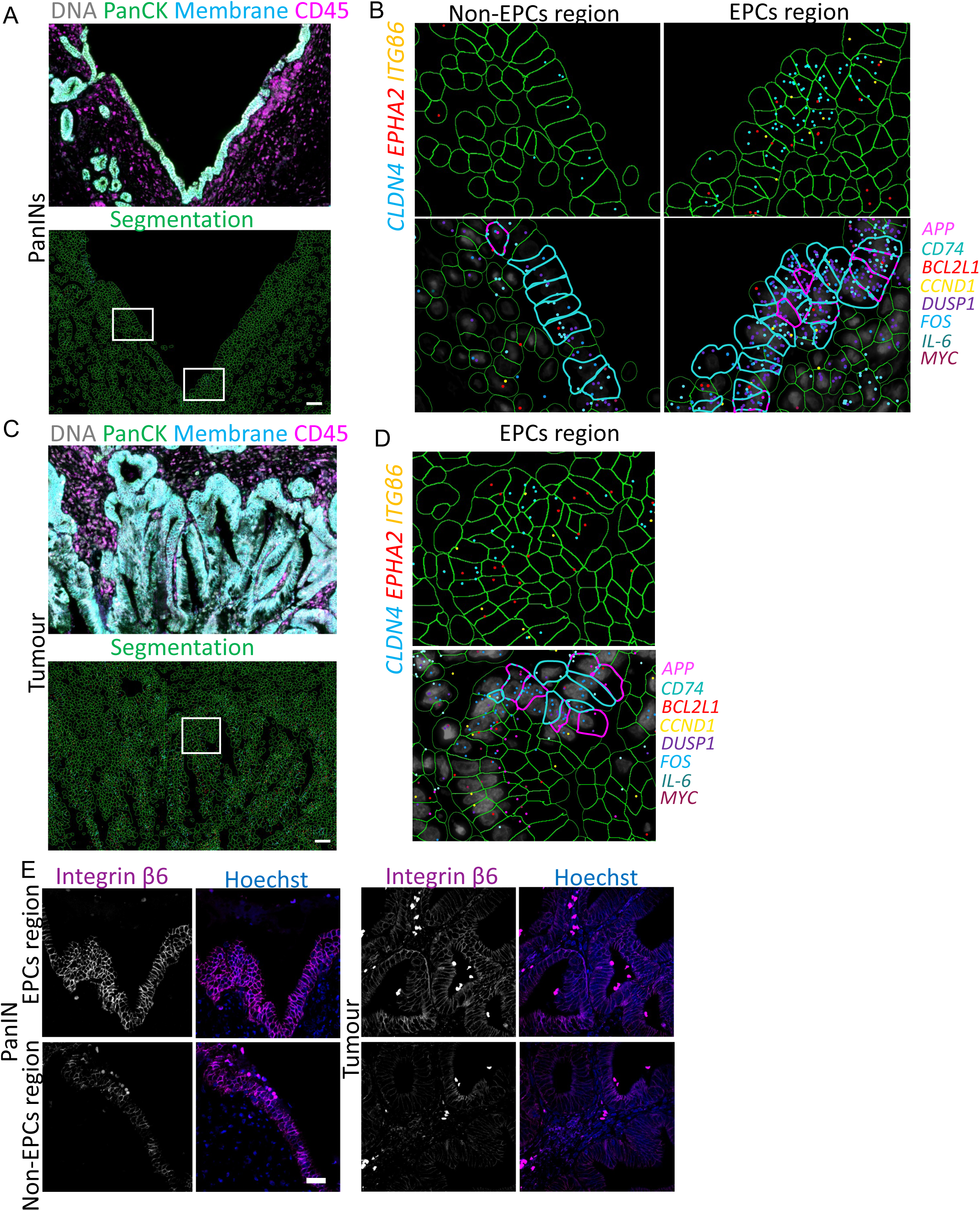
EPCs are present in human PanINs and tumours and show possible interactions with other cells through APP-CD74 crosstalk. (A, C) Representative CosMx images of human pancreatic tissue showing a PanIN (A) and a tumour (C) region stained with Pan-cytokeratin (PanCK), and CD45 antibodies and membrane marker (top) and corresponding cell segmentation (bottom). (B, D) Spatially resolved transcript detection and cell segmentation used to define individual cells. Top panels show dots representing RNA probe detection of *CLDN4*, *EPHA2* and *ITGΒ6* EPC gene markers. In B, two regions have been identified from A: EPCs and non-EPCs regions, containing high or low numbers of EPC markers expressing cells respectively. Bottom panel show the expression of *APP* (pink segmentation), *CD74* (cyan segmentation), *BCL2L1*, *CCND1*, *DUSP1*, *FOS*, *IL6* and *MYC* genes. (E) Representative images of human PanIN (left) and tumour (right) tissues stained with Integrin β6 and Claudin 4 at different timepoints. Scale bar: 50μm.

Our analysis suggests a critical role of CD74 activity in epithelial cell survival via MAPK-ERK and NFκB pathways in epithelial cells in pancreatic cancer development. CD74 inhibition decreases organoid growth/cell survival in HG-PanIN organoids, suggesting CD74 is essential for HG-PanIN survival and growth. To test if APP-CD74 interactions were also present in human tissues, we identified *APP* and *CD74* gene expression in cells and compared levels of expression in EPCs and non-EPC regions in human PanINs and in tumours using spatial transcriptomics. We found some cells expressing EPC markers also expressed *APP* and these cells were surrounded by *CD74* expressing cells, including EPCs and to a lesser extent non-EPCs (Figure 6A-D bottom panels; delineated cells in magenta =*APP* and cyan =*CD74*). To test if the APP-CD74 crosstalk was activating ERK1/ERK2 and NFκB pathways, we interrogated the expression of downstream genes activated in these pathways and present in the HUCP including *FOS, CCND1, DUSP1 and MYC*^27,53^ for ERK1/ERK2 pathway, and *IL6*, *CCND1* and *BCL2L1*^54–56^ for NFκB pathway. We found that expression of these genes was increased in *CD74* expressing cells adjacent to *APP* expressing cells in PanINs and tumours in EPCs regions, while they were poorly expressed in *CD74* expressing cells in non-EPC regions (Figure 6A-D, bottom panels).

In conclusion, our results show the presence of EPCs in human PanINs and tumours. EPCs neighbouring EPCs or non-EPC epithelial cells in PanINs and tumours express APP and CD74 genes, suggesting APP-CD74 ligand-receptor interactions are also relevant in human. Gene expression data suggest ERK1/ERK2 and NFκB pathways are higher in CD74-receiving cells, confirming and extending the results obtained in mouse tissues.

## Discussion

Here, we identify and characterise a novel population of pancreatic cancer stem-like cells: the Early Progression Cells (EPCs). By combining our established mouse model of sporadic tumorigenesis with chronic inflammation, transcriptomic analyses and immunofluorescence imaging, we show that EPCs represent a distinct population of cells that are frequently detected in early precancerous stages (ADM/metaplasia, LG-PanINs) and in low-grade to high-grade PanIN lesions but are less prevalent in tumours. Our results show that approximately 10% of PanINs/GLS contain more than 30% of EPCs and PanINs enriched for EPCs are more proliferative than those with fewer EPCs. *In vitro* sphere formation assays confirmed the higher stemness potential of EPCs compared to other cells in LG-PanINs and HG-PanINs. Recent studies show that 60% of healthy adults carry PanINs in their pancreas^65^, however, only ∼0.05% per year develop pancreatic cancer, suggesting only rare PanINs progress into tumours. The cellular and molecular characteristics of tumour-progressing PanINs remain poorly understood. Our data suggest that PanINs enriched for EPCs progress towards tumours; however, further studies are needed to determine whether EPC^high^ PanINs specifically drive tumorigenesis *in vivo*. Importantly, we demonstrate that EPC signatures increase over timelines to tumorigenesis *in vivo*, and human tumours contain regions enriched for EPC protein and gene markers, strongly suggesting that EPCs play a critical role in tumorigenesis.

Previous studies have identified ‘progenitor-like’ populations essential to drive pancreatic cancer development^13,21^. Like previous reports, we find that EPCs share gene signatures of progenitor-like cells (e.g., p53 pathway, apoptosis, TGFβ wound healing responses). Equally, we show that EPCs are distinct and express gene signatures indicative of immune responses, cell stress responses, survival pathways and remodelling of cell-matrix adhesion, as well as specific gene and protein markers of dedifferentiated epithelial cells (e.g., Integrin β6, Claudin 4, Sca-1 and EphA2)^57,58^. Our transcriptomic trajectory and pseudotime analyses reveal that EPCs are heterogenous in cell fate and behave as putative ancestral cells, giving rise to stem-like cells, neoplastic/metaplastic and regenerating acinar cells. We identified a second cluster of stem-like cells (cluster 5) that pseudotime analysis positioned as possibly derived from EPCs and express progenitor and stem gene signatures, including embryonic and pancreatic cancer stem cell signatures. Our analyses suggest that cluster 5 gene signatures align to some extent with previously reported signatures of pancreatic cancer progenitor-like/malignant cells in PanINs^13,21^ (Figure S4).

EPC identity and heterogeneity in cell fate likely reflect our unique approach to modelling sporadic pancreatic cancer in the mouse *in vivo*. In contrast to previously published studies^12,13,21^, we induce oncogenic KrasG12D mutations in low numbers of epithelial cells in adult tissues^9,10^, enabling us to ask how rare genetically mutant cells give rise to tumours. Using this approach, we have revealed the importance of cell-cell interactions between mutant and non-mutant epithelial cells in pancreas health and early pancreatic cancer^9,10^. Here, we use single cell/nuclei RNA sequencing of fixed tissues, which we believe has allowed us to better capture exocrine populations^59^ and infrequent events that would otherwise be underrepresented using protocols that involve cell dissociation. However, a limitation of this approach is our inability to robustly determine gene signatures of RFP-labelled KrasG12D-expressing cells in the transcriptomic data. RFP probes failed to identify the number of cells consistent with the percentage of RFP positive cells expected in tissues, probably due to the low expression of RFP transcripts in the cytoplasm and nuclei of labelled cells. Therefore, we are unable to directly rule out the contribution of non-mutant cells to EPC populations in the transcriptomic data. EPCs (cluster 6) significantly upregulate Kras driven pathways compared to other epithelial cell clusters, including Hallmark Kras signalling up, Reactome and Wikipathways MAPK signaling or GOBP ERK1 and ERK2 cascade, suggesting they are Kras mutant cells; however, whether EPC cell fate correlates with Kras mutational status requires further investigation. Future studies will delineate the mechanisms controlling EPC transitions during disease progression.

Our transcriptomic data indicate that EPCs upregulate stress-related pathways, and exhibit high expression of cytoprotective and stress-adaptive factors, including Nupr1 and Clusterin (*Clu*), both of which are known to promote survival under conditions of oxidative and genotoxic stress^44,66^. Nupr1 has been shown to play an essential role in pancreatic tumorigenesis, protecting cells from stress-induced death by inhibiting apoptosis through a pathway dependent on transcription factor RelB and immediate early response 3 (IER3)^44^. Our results showed Nupr1 to be upregulated in acinar cell clusters, consistent with an upregulation of Nupr1 in acinar cells undergoing chronic pancreatitis^67^, and in cluster 6 (EPCs) and cluster 13, which sits very closely to cluster 6. Our results showed that Nupr1 is critical for the progression of PanIN/tumour organoid growth, with very few organoids surviving Nupr1 inhibition. Consistent with our results, previous reports showed that ZZW-115 prevented the development of PanINs in KC mice, triggering cell death in KRasG12D-expressing cells specifically^68^. Clusterin is expressed in acinar cells undergoing injury-like reprogramming^60^ and in pancreatitis-induced ADM, where it protects cells from apoptosis^69^, while Clusterin inhibition in pancreatic cancer represses proliferation inducing senescence^70^. Here we found Clusterin to be highly upregulated in clusters 5 and 6, coinciding with the most stem-like populations. Notably, while we see intrinsic apoptotic signalling pathways enriched, pathways involved in the negative regulation of apoptosis are also significantly upregulated, indicating that apoptotic signals are actively restrained. Together, these data support a model in which EPCs exist in a stress-adapted, apoptosis-poised state that enables survival and progression during early tumorigenesis. Further research needs to be done to functionally confirm the ability of EPCs to survive stress.

We found that APP-CD74 is the most upregulated ligand-receptor interaction between EPCs and neighbouring exocrine cells. CD74 is a type II transmembrane glycoprotein associated with the major histocompatibility complex class II (MHC II) and as such, CD74 is best described in immune cells. Nonetheless, CD74 has been linked to epithelial tumours including pancreatic cancer (95% of pancreatic carcinomas)^71^ and is associated with perineural invasion and poor prognosis^72^. While CD74 is rarely expressed in acinar cells, it is highly expressed in tissues with pancreatitis and PanINs, increasing in expression with grade progression^71^. Consistent with our results, knockdown of CD74 in pancreatic cancer cell lines leads to decreased cell proliferation, migration, and invasion^73^. The role of APP in pancreatic tissue injury and pancreatic cancer and the importance of APP-CD74 interactions in epithelial tissues is not well understood. Here we report that APP-CD74 interactions between epithelial cells modulate neoplastic cell growth. Our results illustrate an activation of ERK1/ERK2 pathway coupled with increased nuclear pERK in CD74 positive epithelial cells in PanINs and tumours. We also show that NFκB pathway is activated in CD74 expressing cells and increased NFκB in the nuclei of CD74 positive epithelial cells, suggesting that both pathways are activated downstream of CD74 activation. The PI3K/AKT pathway was not significantly enriched and was even reduced in tumour CD74 positive cells at the gene level, suggesting PI3K pathway is not required in our model. We therefore hypothesise that in the context of pancreatic cancer initiation, CD74 signalling may preferentially activate ERK/NFκB pathways while PI3K/AKT signalling is not engaged. Future studies are required to mechanistically understand APP-CD74 downstream signalling in EPCs and in the context of EPC-containing PanINs.

Pancreatic cancer is very often diagnosed at late and incurable stages. A better understanding of the early stages of disease is critical to the discovery of new and improved strategies to prevent, detect and intercept disease earlier. Chronic pancreatitis is strongly associated with increased risk of pancreatic cancer, particularly within 2 years of pancreatitis diagnosis^61^. Our results position EPCs as critical players in pancreatic cancer initiation and progression in a model of KrasG12D-driven disease coupled with chronic pancreatitis. Further study of these cells could lead to the development of tools to detect patients at high risk of developing pancreatic cancer and new therapeutics to prevent cancer development.

## Acknowledgments

The authors thank A. Marchbank and colleagues at School of Biosciences Genome Hub (Cardiff University) and Wales Gene Park for technical support with RNA sequencing experiments. The authors thank colleagues at School of Biosciences Animal Facility (Cardiff University) for their support with the animal work. Finally, this work was funded by Pancreatic Cancer UK Career Foundation Fellowship to BSB CFF2022_Salvador, Pancreatic Cancer UK Research Innovation Fund to BSB and CH RIF2023-24_Salvador and philanthropic donation to CH lab from Stephen A. Hopkins family. The authors thank the Pancreatic Cancer Research Fund Tissue Bank for providing the human samples used in this study.

## M&M

### Mouse strain and treatment protocol

Animals were housed in conventional pathogen-free animal facilities and experiments were conducted in accordance with UK Home Office regulations (ASPA 1986 and EU Directive 2010) under the guidelines of Cardiff University Animal Welfare and Ethics Committee. Pdx1-CreERT; LSL-KrasG12D/+^62^; Rosa26 LSL-tdRFP^63^ male and female 6-to 8-week-old mice were induced by one intraperitoneal injection of 1μg/40g bodyweight tamoxifen in corn oil as previously described^9,10^. Five weeks later, mice were treated with five doses of 0.2mg/kg of caerulein (Sigma-Aldrich) a week for 3 weeks for ADM timepoint, or 6 weeks to induce PanINs/tumours. Mice were subsequently aged to defined timepoints representative of LG-/HG-PanINs and tumour.

### Tissue staining

Pancreatic tissue was harvested at the indicated experimental timepoints and immersion-fixed overnight at 4°C in 10% neutral buffered formalin. Following fixation, each pancreas was divided into two portions (head-body and body-tail) and processed for either paraffin or OCT matrix embedding. Pancreatic human tissues containing PanINs/tumours were annotated and provided by Pancreatic Cancer Research Fund Tissue Bank and with ethics approved by their Tissue Access Committee (TAC) and Operations Group. Formalin-fixed paraffin-embedded (FFPE) mouse tissues were sectioned at 7 μm thickness and processed for immunofluorescence using the antibodies listed in Table 1.

**Table 1.**
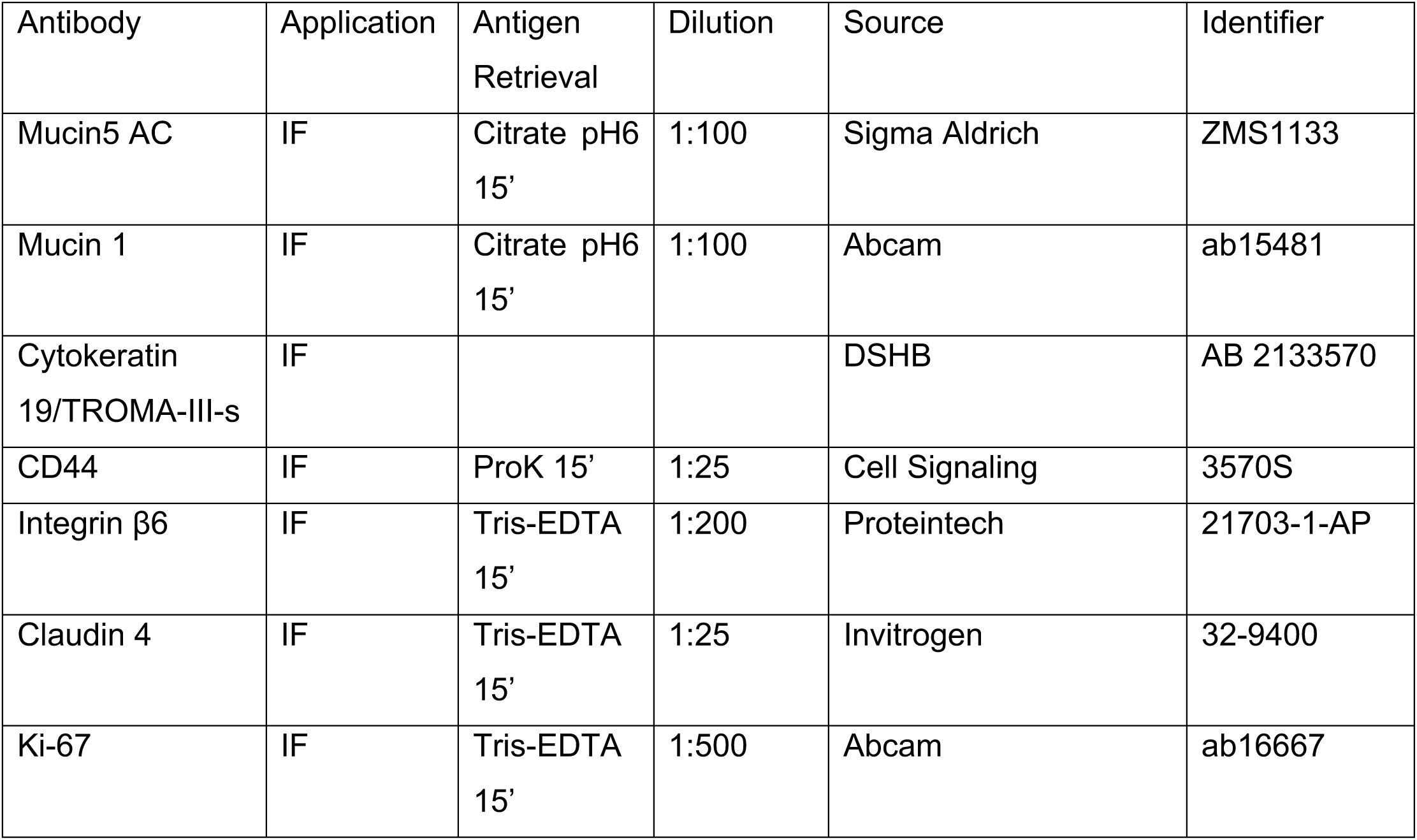
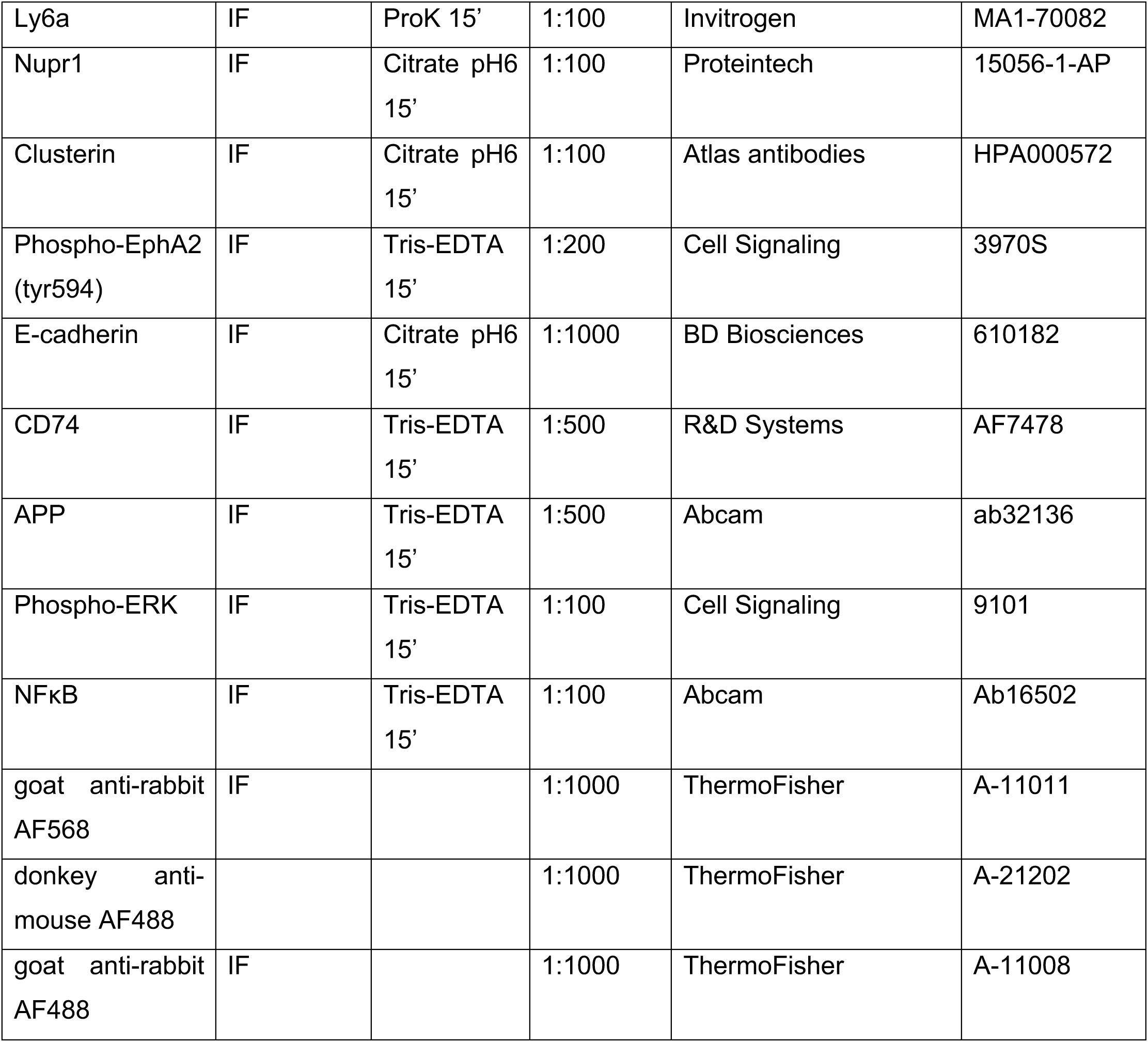

For immunostaining, FFPE sections were deparaffinised and rehydrated before antigen retrieval. Depending on the primary antibody, antigen retrieval was performed either by incubation with 20μg/ml Proteinase K in TBS-T for 15 min at 37°C, or by heat-mediated retrieval in citrate buffer (pH 6) or Tris-EDTA buffer (pH 9) for 15 min, as indicated in Table 1. Sections were subsequently blocked with 5% normal goat serum (NGS) in TBS-T for 30-60 min at room temperature. Primary antibodies were diluted in 5% NGS/TBS-T at the concentrations specified in Table 1 and applied overnight at 4°C. Following washing, sections were incubated with the appropriate Alexa Fluor-conjugated secondary antibodies for 60 min at room temperature and counterstained with Hoechst (Thermo Fisher Scientific). Slides were mounted using Mowiol (Sigma-Aldrich).

For each mouse, the endogenous RFP-positive area was quantified across five 10 μm formalin-fixed OCT-embedded frozen sections, with a minimum interval of 20 μm between consecutive sections. Fluorescence imaging was performed using a Zeiss LSM710 confocal microscope.

PanIN lesions were identified in FFPE sections following Alcian blue staining, as previously described^9^.

### Image analysis

Endogenous RFP levels were quantified as previously described^10^. Using Fiji (RRID:SCR_002285) software, positive areas were thresholded using RFP fluorescence. Tissue autofluorescence was used to determine total tissue area.

Integrin β6, Claudin 4, Cytokeratin 8, Sca-1, Nupr1, Clu and pEphA2 positive cells per PanIN/GLS were assessed manually by quantifying cells with staining in the membrane around the whole cell (Integrin β6, Claudin 4, Cytokeratin 8, Sca-1, Clu and EpCAM), apical membrane (pEphA2) or high nuclear (Nupr1) staining. Ki-67 nuclear staining was quantified as positive. Finally, pERK and NFκB nuclear fluorescence intensity was quantified using Fiji and only cells with cytoplasmic staining were considered positive for CD74.

### Single Cell/Nuclei RNA sequencing

Single cell/nuclei RNA sequencing was performed using the GEM FLEX 10X Genomics protocol on FFPE sections, following manufacturer’s instructions. Sixteen single-cell RNA-seq libraries were generated, spanning six timepoints of pancreatic disease progression: 7 days post-induction (7_dpi; n = 2 libraries), 35 days post-induction (35_dpi; n = 2), acinar-to-ductal metaplasia (ADM; n = 3), low-grade pancreatic intraepithelial neoplasia (LG-PanINs; n = 3), high-grade pancreatic intraepithelial neoplasia (HG-PanINs; n = 3), and tumour (Tumours; n = 3). We QC filtered the samples and per-sample Seurat objects were integrated using Seurat’s canonical correlation analysis (CCA)-based integration workflow (Seurat v4/v5) and the top 2,000 variable features shared across samples. The integrated assay, counting on 149,712 cells, was scaled, and clustering was performed on the first 30 principal components using a shared nearest-neighbour graph followed by Louvain clustering (resolution = 0.5). Uniform Manifold Approximation and Projection (UMAP) was used for two-dimensional visualisation (dims = 1:30). Clusters from the integrated dataset were manually annotated to cell types based on marker gene expression previously published^7,8^, yielding the following populations: Acinar cells (*Cpa1*), Duct-like cells (*Krt19*), Metaplastic cells (*Krt19*, *Clu*), Neoplastic cells (*Arg2*, *Mucin5ac*, *Tff1*), Fibroblasts (*Col1a1*), Macrophages (*Cd68*), T cells (*Cd3d*, *Cd3e*, *Cd3g*), B cells (*Cd19*), Endothelial cells (*Pecam1*), Neuroendocrine cells (*Resp18*), and Pericytes (*Rgs5*).

To focus on the epithelial exocrine compartment, cells annotated as Metaplastic, Duct-like, Neoplastic and Acinar cells were subset from the integrated object and re-clustered independently (resolution = 0.2), with UMAP re-computed on the same dimensions.

Cell cycle phase was scored for each cell using Seurat’s built-in S-phase and G2/M-phase gene sets. One cluster (cluster 8) within the epithelial subset was identified as a discrete population of actively cycling cells based on Phase distribution and was removed to eliminate cell-cycle-driven transcriptional variation as a confounding factor in downstream analyses.

Following removal of the cycling cluster, the epithelial subset was re-scaled and re-clustered from scratch (resolution = 0.3), and UMAP re-computed (dims = 1:20) with the final object used for all subsequent analyses. Cluster 6 of this object was identified as a the PCDC population based on marker expression and downstream trajectory analysis.

Cluster-specific marker genes were identified for each cluster (one cluster vs. all others). Genes were ranked by average log2 fold-change.

To characterise the transcriptional identity of cluster 6, differential expression between cluster 6 and all other clusters within the epithelial subset was calculated and gene set enrichment analysis (GSEA) was performed against publicly available gene signatures. Genes were ranked by average log2 fold-change and tested for enrichment against Hallmark, KEGG_legacy, Reactome and Gene Ontology Biological Process (GOBP) gene sets. Pathways with Benjamini-Hochberg adjusted p < 0.05 were considered significant.

Pseudotime trajectory analysis was performed using Monocle3. The Seurat object (exocrine epithelial subset, cycling cells excluded) was converted to a Monocle3 preserving raw counts, existing UMAP coordinates, and all metadata (cluster identity, timepoint). Pseudotime was calculated with cluster 6 specified as the trajectory root, consistent with its identification as a putative stem/metaplastic population. Pseudotime was validated against the independently defined biological timepoints (7_dpi to 35_dpi to ADM to LG-PanINs to HG-PanINs to Tumour). Gene modules co-varying with pseudotime were identified and aggregated by cluster and by timepoint for visualisation.

Cell-cell communication was inferred using CellChat^41^ (mouse ligand-receptor database). Log-normalised expression data and cluster labels from the epithelial subset were used to construct a CellChat object and over-expressed genes and ligand-receptor interactions were identified, communication probabilities were computed, and interactions supported by fewer than 10 cells were filtered. Pathway-level communication probabilities and the aggregated cell-cell communication network were computed. Ligand-receptor pairs with cluster 6 as the signal source were extracted. The App-Cd74 ligand-receptor pair, with cluster 6 as the source (App) and the remaining clusters as receivers (Cd74), was identified as the most strongly upregulated interaction by this analysis.

To evaluate transcriptional activity of canonical signalling pathways downstream of CD74, Cd74 (log-normalised expression) was extracted directly from the Seurat object’s per-sample expression layers. Cells were stratified into Cd74-high and Cd74-low groups by dividing cells into expression quartiles independently within each timepoint (to account for differences in the overall distribution of Cd74 expression across timepoints), retaining the top 25% (Cd74-high) and bottom 25% (Cd74-low) of cells per timepoint for downstream comparison; the middle 50% of cells were excluded from group comparisons. NF-κB, PI3K/AKT, and ERK1/ERK2 signalling per-cell module scores were calculated using Seurat’s ‘AddModuleScore()’. Gene sets were retrieved from MSigDB: HALLMARK_TNFA_SIGNALING_VIA_NFΚB (NF-κB), HALLMARK_PI3K_AKT_MTOR_SIGNALING (PI3K/AKT), and GOBP_ERK1_AND_ERK2_CASCADE (Gene Ontology, GO:0070371); (ii) GOBP_POSITIVE_REGULATION_OF_ERK1_AND_ERK2_CASCADE and GOBP_NEGATIVE_REGULATION_OF_ERK1_AND_ERK2_CASCADE (GO:0070374 and GO:0070372 respectively). Pathway module scores were compared between Cd74-high and Cd74-low cells using the Wilcoxon rank-sum test, performed separately at each timepoint. Spearman rank correlation was used to assess the continuous relationship between Cd74 expression level and each pathway module score.

### Sphere formation assay

Pancreas tissues were harvested LG-PanINs or HG-PanINs timepoints. Pancreas tissues were digested as previously described^64^. Using an EGTA-based buffer the pancreas from each mouse was inflated and slightly digested with collagenase. The tissue was chopped and further digested using a calcium-based buffer with collagenase. Single cell digested tissue was stained with anti-Sca-1/Ly6a (Rat) and anti-Mucin5ac (Rabbit) primary antibodies (1μl/ml) for 20 min in EGTA-based buffer at 4C. Then cells were washed twice and stained with FITC anti-Rat antibody for 15 min in EGTA-based buffer at 4°C. Cells were then labelled with MACS anti-FITC magnetic beads and separated using magnetic cell isolation columns following manufacturer’s instructions. Unlabelled recovered cells were then stained with FITC anti-Rabbit secondary antibody and separated again following the protocol described before. Sca-1 negative cells were treated to the same protocol as Sca-1 positive cells but without secondary antibody. Recovered cells were then transferred to an Ultra-low Attachment 24MW plate, with 5.000 cells per well (3 wells per condition (Sca-1 positive or Sca-1 negative Mucin5ac positive) and cultured in DMEM/F12 (Gibco) with 1% Pen/Strep (Gibco), 1% Fungizone (Gibco), 1% L-Glutamine (Gibco), 1% B-27 (Gibco) and 0.01μg/μl FGFb (Preprotech)^65^. Cells were cultured at 37°C 5% CO2 for 14 days adding additional culture medium if necessary. Spheres were counted manually under a brightfield microscope and the diameter measured using Fiji (RRID:SCR_002285).

### Organoid experiments

Mouse pancreatic organoids from LG-PanINs, HG-PanINs and Tumours timepoints were generated and validated following previously published protocols^66^. Organoids were seeded at equal concentration and 24h later treated with 5μM ZZW-115 or DMSO for Nupr1 inhibition experiments, or 5μg/ml Milatuzumab or IgG in equal concentration for CD74 inhibition experiments. Organoids were monitored for 45-150h in an Incucyte S3 (Sartorius) Organoid module for live cell imaging.

### Spatial transcriptomics

Pancreatic human tissues containing PanINs/tumours were annotated and provided by Pancreatic Cancer Research Fund Tissue Bank. FFPE tissue sections were prepared for spatial transcriptomic experiments according to the manufacturer’s recommendations (Bruker) and mounted onto CosMx-compatible slides. Tissue sections were processed for RNA detection using the CosMx Human RNA assay, including tissue preparation, permeabilisation, hybridisation and signal amplification steps using the Human Universal Cell Characterisation RNA panel as specified by the manufacturer. Following probe hybridisation and signal amplification, transcript detection and high-resolution imaging were performed using the CosMx Spatial Molecular Imager. Tissue segmentation and transcript detection was performed using the AtoMx Spatial Informatics Platform.

### Statistical tests

Statistical analyses were performed using GraphPad Prism 10 and 11 or R studio software. Normally distributed data, as determined by the Shapiro-Wilke test were analysed using unpaired Student’s t tests with Welch correction. Data that were not normally distributed were compared using non-parametric Mann Whitney U two-tailed t tests. A p value of < 0.05 was taken as significant. No statistical method was used to pre-determine sample size. Definition of n defined in the figure legends.

