## Supplementary material for "Early Progression Cells mark a stem-like state associated with pancreatic cancer initiation and progression": Suplemental figures

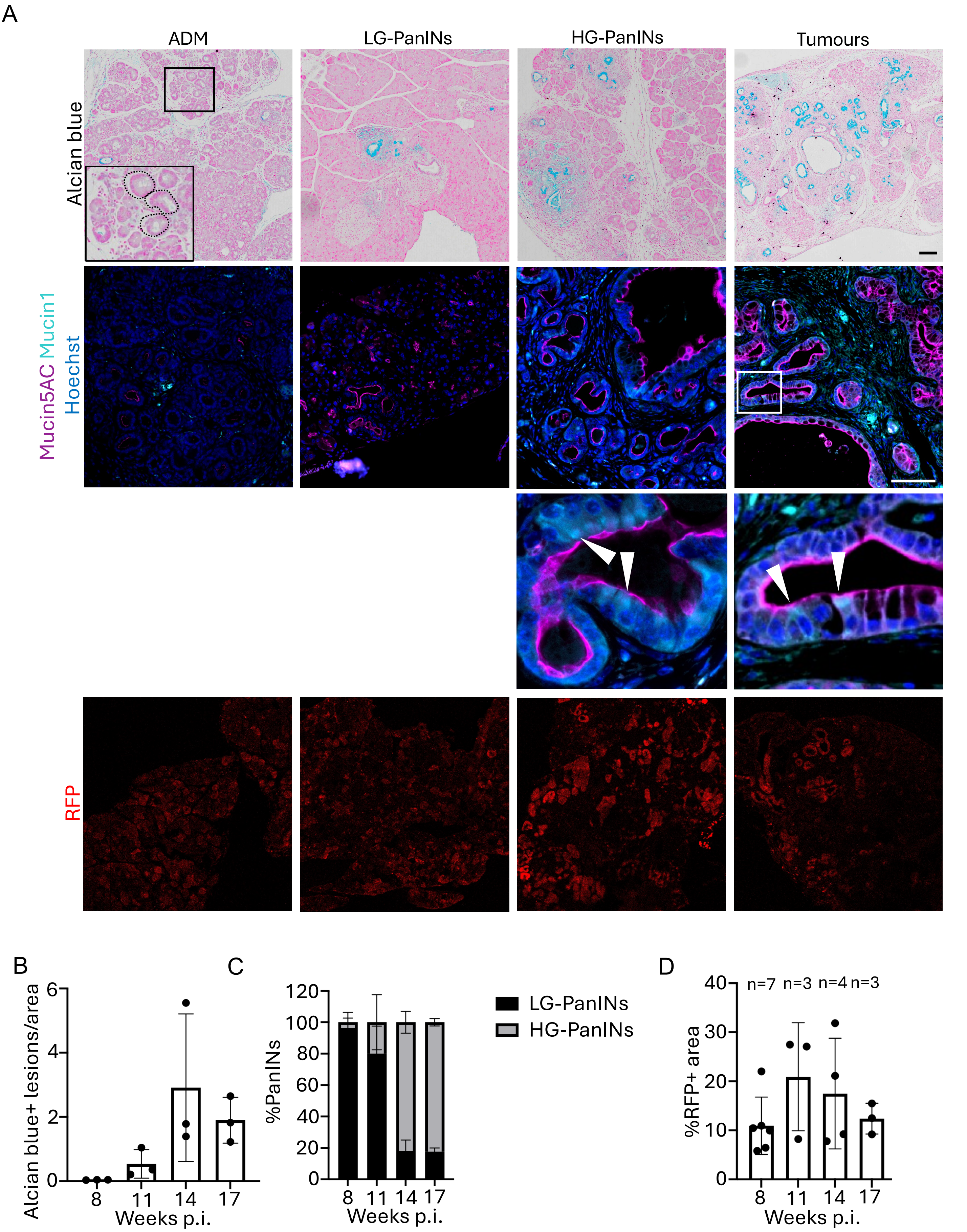
A
ADM
LG-PanINs
HG-PanINs
Tumours
Alcian blue
Mucin5AC Mucin1
Hoechst
RFP
B
Alcian blue+ lesions/area
0
2
4
6
8
11
14
17
Weeks p.i.
C
%PanINs
0
20
40
60
80
100
120
8
11
14
17
Weeks p.i.
LG-PanINs
HG-PanINs
D
%RFP+ area
0
10
20
30
40
n=7
n=3
n=4
n=3
8
11
14
17
Weeks p.i.

**Sup Figure 1**

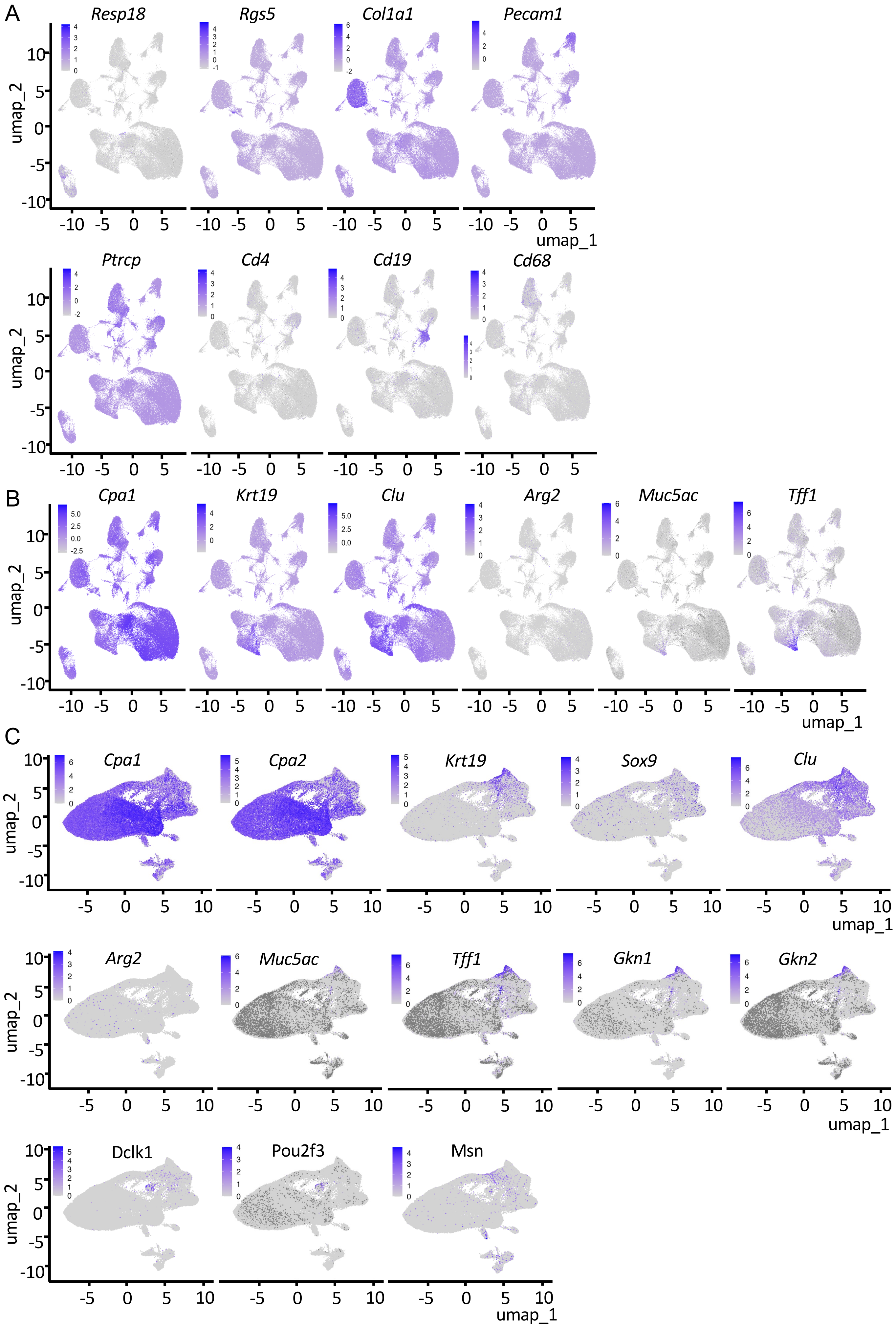
A
Resp18
Rgs5
Col1a1
Pecam1
Ptrcp
Cd4
Cd19
Cd68
B
Cpa1
Krt19
Clu
Arg2
Muc5ac
Tff1
C
Cpa1
Cpa2
Krt19
Sox9
Clu
Arg2
Muc5ac
Tff1
Gkn1
Gkn2
Dclk1
Pou2f3
Msn
umap_1
umap_2

**Sup Figure 2**

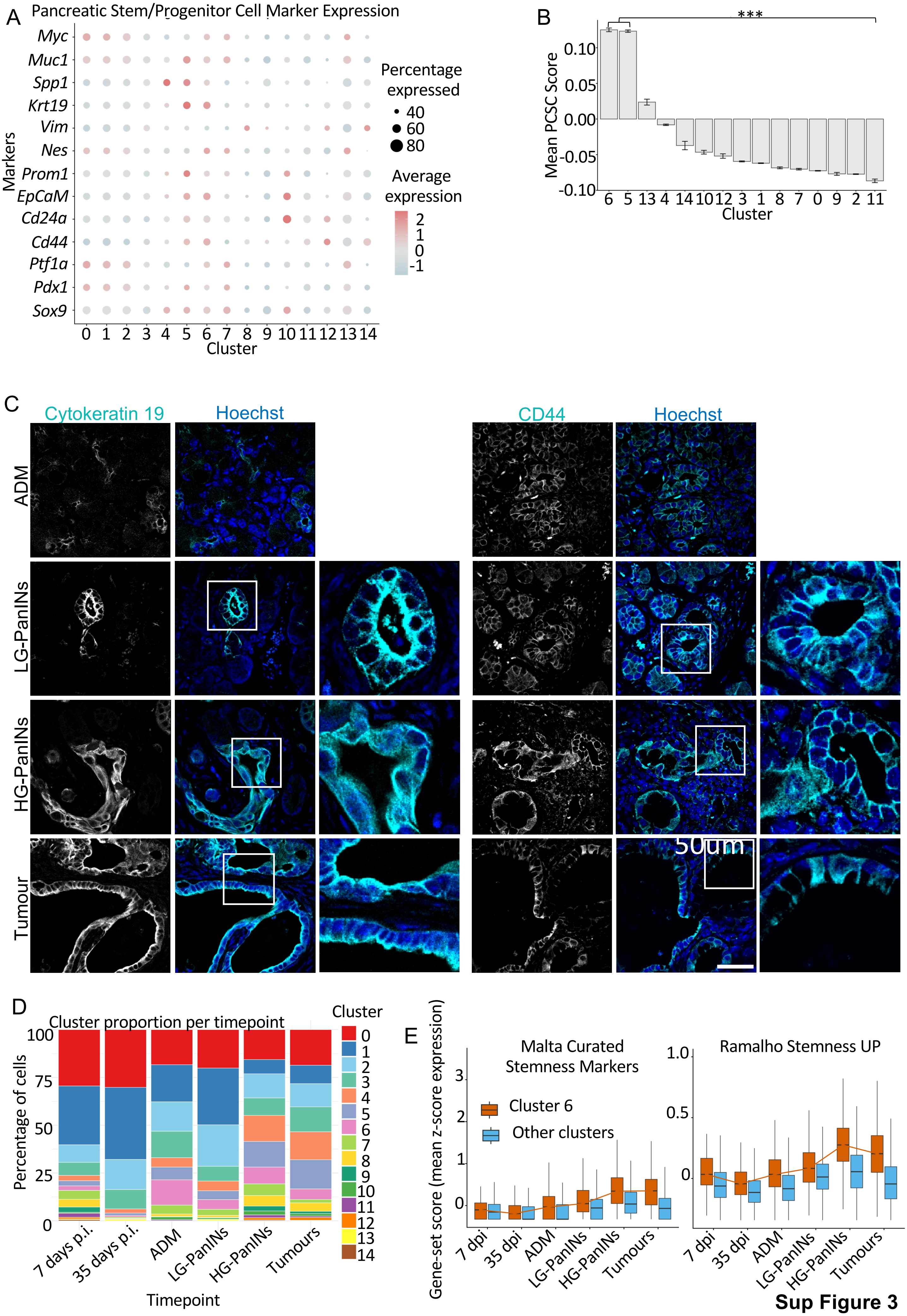
A
Pancreatic Stem/Progenitor Cell Marker Expression
Myc
Muc1
Spp1
Krt19
Vim
Nes
Prom1
EpCaM
Cd24a
Cd44
Ptf1a
Pdx1
Sox9
Markers
0 1 2 3 4 5 6 7 8 9 10 11 12 13 14
Cluster
Percentage expressed
40
60
80
Average expression
2
1
0
-1
B
***
Mean PCSC Score
0.10
0.05
0.00
-0.05
-0.10
6 5 13 4 14 10 12 3 1 8 7 0 9 2 11
Cluster
C
Cytokeratin 19
Hoechst
CD44
Hoechst
ADM
LG-PanINs
HG-PanINs
Tumour
50um
D
Cluster proportion per timepoint
Percentage of cells
100
75
50
25
0
7 days p.i.
35 days p.i.
ADM
LG-PanINs
HG-PanINs
Tumours
Timepoint
Cluster
E
Gene-set score (mean z-score expression)
Malta Curated Stemness Markers
Cluster 6
Other clusters
3
2
1
0
7 dpi
35 dpi
ADM
LG-PanINs
HG-PanINs
Tumours
Ramalho Stemness UP
1.0
0.5
0

**Sup Figure 3**

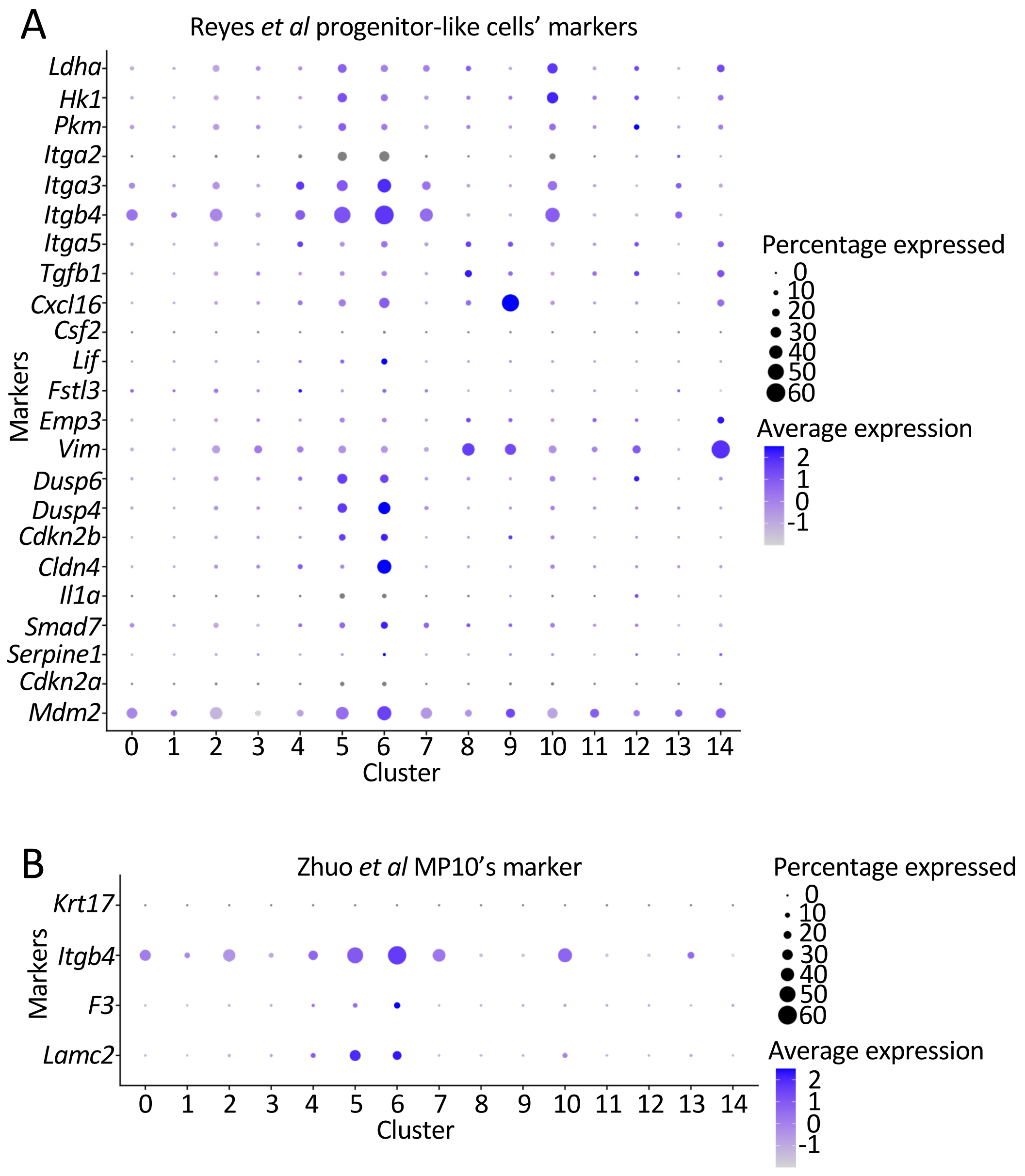

A
Reyes *et al* progenitor-like cells' markers
Ldha
Hk1
Pkm
Itga2
Itga3
Itgb4
Itga5
Tgfb1
Cxcl16
Csf2
Lif
Fstl3
Emp3
Vim
Dusp6
Dusp4
Cdkn2b
Cldn4
Il1a
Smad7
Serpine1
Cdkn2a
Mdm2
Markers
0 1 2 3 4 5 6 7 8 9 10 11 12 13 14
Cluster
Percentage expressed
0
10
20
30
40
50
60
Average expression
2
1
0
-1
B
Zhuo *et al* MP10's marker
Krt17
Itgb4
F3
Lamc2
Markers
0 1 2 3 4 5 6 7 8 9 10 11 12 13 14
Cluster
Percentage expressed
0
10
20
30
40
50
60
Average expression
2
1
0
-1

**Sup Figure 4**

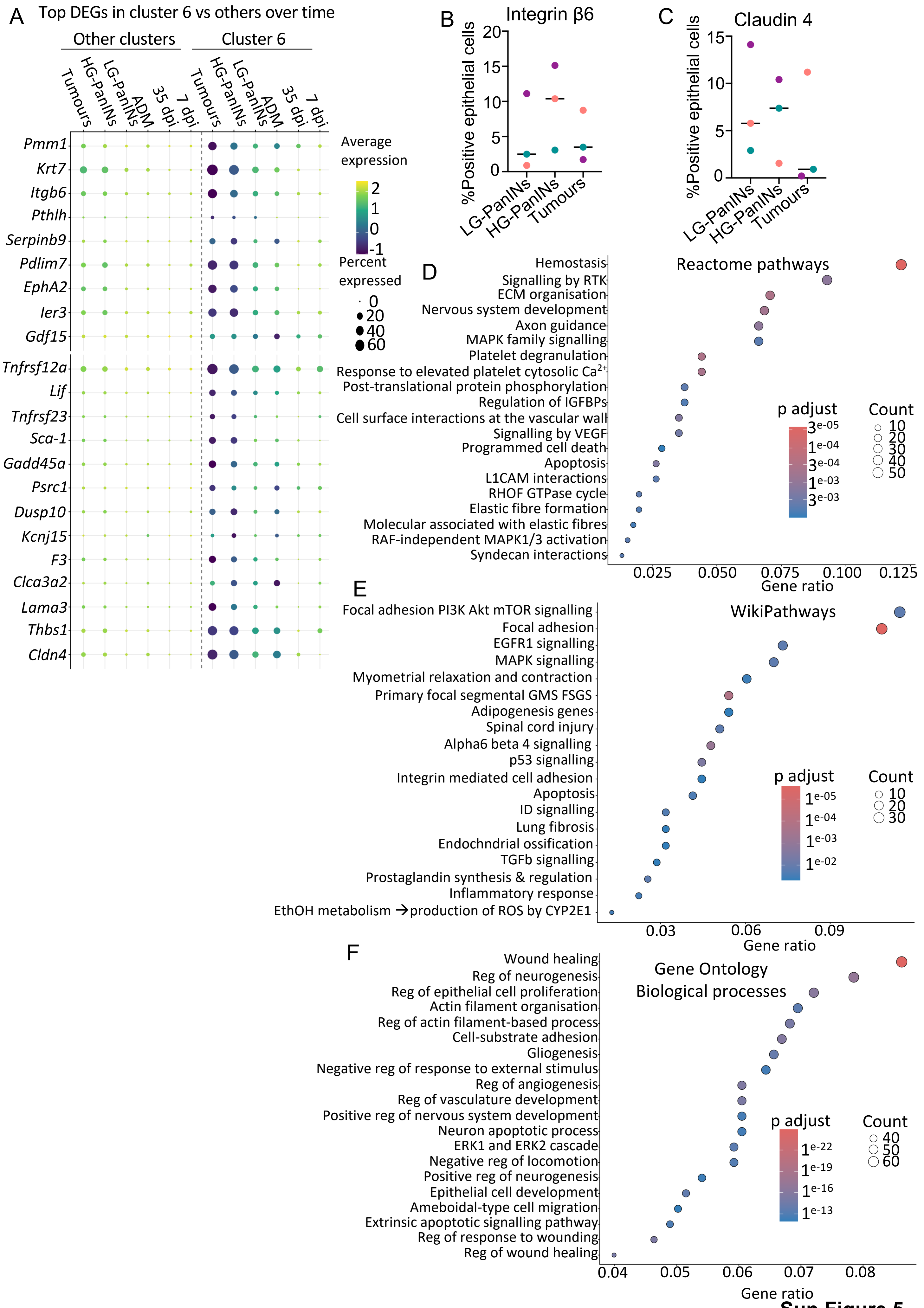
A
Top DEGs in cluster 6 vs others over time
Other clusters
Cluster 6
Tumours
HG-PanINs
LG-PanINs
ADM
35 dpi
7 dpi
Pmm1
Krt7
Itgb6
Pthlh
Serpinb9
Pdlim7
EphA2
Ier3
Gdf15
Tnfrsf12a
Lif
Tnfrsf23
Sca-1
Gadd45a
Psrc1
Dusp10
Kcnj15
F3
Clca3a2
Lama3
Thbs1
Cldn4
Average expression
2
1
0
-1
Percent expressed
0
20
40
60
B
Integrin β6
%Positive epithelial cells
20
15
10
5
0
LG-PanINs
HG-PanINs
Tumours
C
Claudin 4
%Positive epithelial cells
15
10
5
0
LG-PanINs
HG-PanINs
Tumours
D
Reactome pathways
Hemostasis
Signalling by RTK
ECM organisation
Nervous system development
Axon guidance
MAPK family signalling
Platelet degranulation
Response to elevated platelet cytosolic Ca2+
Post-translational protein phosphorylation
Regulation of IGFBPs
Cell surface interactions at the vascular wall
Signalling by VEGF
Programmed cell death
Apoptosis
L1CAM interactions
RHOF GTPase cycle
Elastic fibre formation
Molecular associated with elastic fibres
RAF-independent MAPK1/3 activation
Syndecan interactions
p adjust
3e-05
1e-04
3e-04
1e-03
3e-03
Count
10
20
30
40
50
0.025
0.050
0.075
0.100
0.125
Gene ratio
E
WikiPathways
Focal adhesion PI3K Akt mTOR signalling
Focal adhesion
EGFR1 signalling
MAPK signalling
Myometrial relaxation and contraction
Primary focal segmental GMS FSGS
Adipogenesis genes
Spinal cord injury
Alpha6 beta 4 signalling
p53 signalling
Integrin mediated cell adhesion
Apoptosis
ID signalling
Lung fibrosis
Endochndrial ossification
TGFb signalling
Prostaglandin synthesis & regulation
Inflammatory response
EthOH metabolism →production of ROS by CYP2E1
p adjust
1e-05
1e-04
1e-03
1e-02
Count
10
20
30
0.03
0.06
0.09
Gene ratio
F
Gene Ontology
Biological processes
Wound healing
Reg of neurogenesis
Reg of epithelial cell proliferation
Actin filament organisation
Reg of actin filament-based process
Cell-substrate adhesion
Gliogenesis
Negative reg of response to external stimulus
Reg of angiogenesis
Reg of vasculature development
Positive reg of nervous system development
Neuron apoptotic process
ERK1 and ERK2 cascade
Negative reg of locomotion
Positive reg of neurogenesis
Epithelial cell development
Ameboidal-type cell migration
Extrinsic apoptotic signalling pathway
Reg of response to wounding
Reg of wound healing
p adjust
1e-22
1e-19
1e-16
1e-13
Count
40
50
60
0.04
0.05
0.06
0.07
0.08
Gene ratio

**Sup Figure 5**

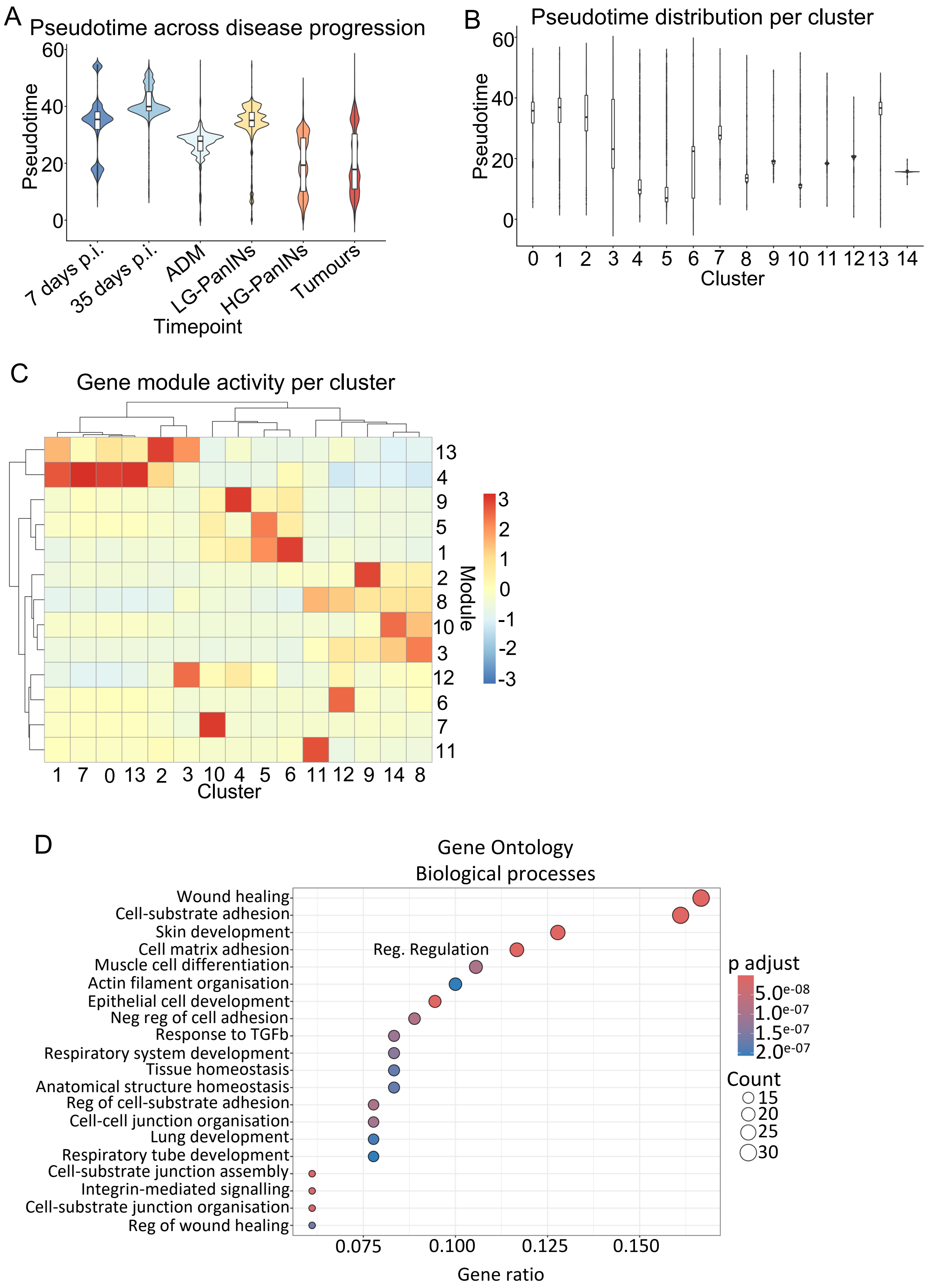
A
Pseudotime across disease progression
Pseudotime
Timepoint
7 days p.i.
35 days p.i.
ADM
LG-PanINs
HG-PanINs
Tumours
B
Pseudotime distribution per cluster
Pseudotime
Cluster
C
Gene module activity per cluster
Module
Cluster
D
Gene Ontology
Biological processes
Wound healing
Cell-substrate adhesion
Skin development
Cell matrix adhesion
Reg. Regulation
Muscle cell differentiation
Actin filament organisation
Epithelial cell development
Neg reg of cell adhesion
Response to TGFb
Respiratory system development
Tissue homeostasis
Anatomical structure homeostasis
Reg of cell-substrate adhesion
Cell-cell junction organisation
Lung development
Respiratory tube development
Cell-substrate junction assembly
Integrin-mediated signalling
Cell-substrate junction organisation
Reg of wound healing
Gene ratio
p adjust
5.0e-08
1.0e-07
1.5e-07
2.0e-07
Count
15
20
25
30

**Sup Figure 6**

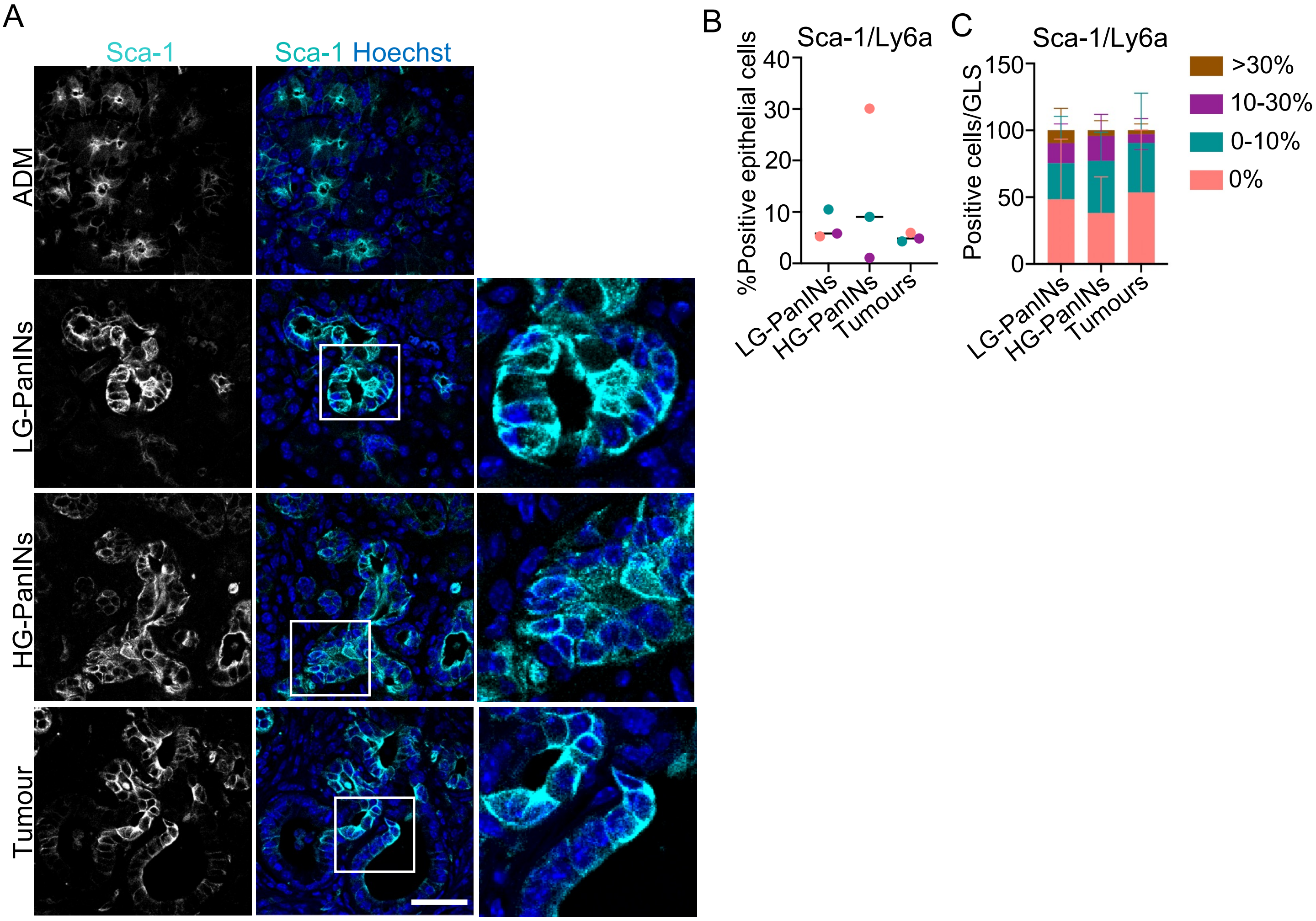
A
Sca-1
Sca-1 Hoechst
ADM
LG-PanINs
HG-PanINs
Tumour
B
Sca-1/Ly6a
%Positive epithelial cells
40
30
20
10
0
LG-PanINs
HG-PanINs
Tumours
C
Sca-1/Ly6a
Positive cells/GLS
150
100
50
0
LG-PanINs
HG-PanINs
Tumours
>30%
10-30%
0-10%
0%

**Sup Figure 7**

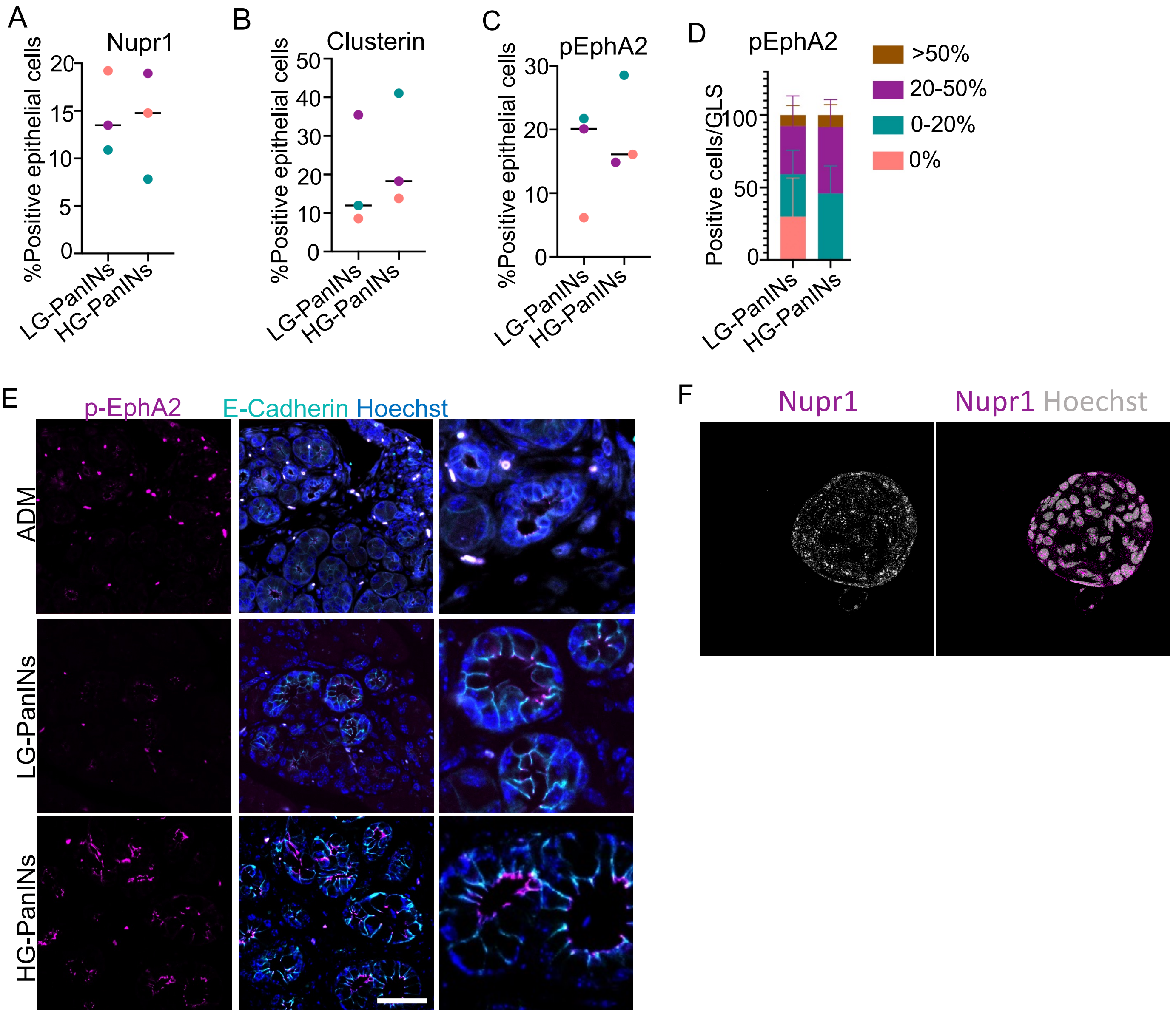
A
Nupr1
%Positive epithelial cells
20
15
10
5
0
LG-PanINs
HG-PanINs
B
Clusterin
%Positive epithelial cells
50
40
30
20
10
0
LG-PanINs
HG-PanINs
C
pEphA2
%Positive epithelial cells
30
20
10
0
LG-PanINs
HG-PanINs
D
pEphA2
Positive cells/GLS
100
50
0
LG-PanINs
HG-PanINs
>50%
20-50%
0-20%
0%
E
p-EphA2
E-Cadherin Hoechst
ADM
LG-PanINs
HG-PanINs
F
Nupr1
Nupr1 Hoechst

**Sup Figure 8**

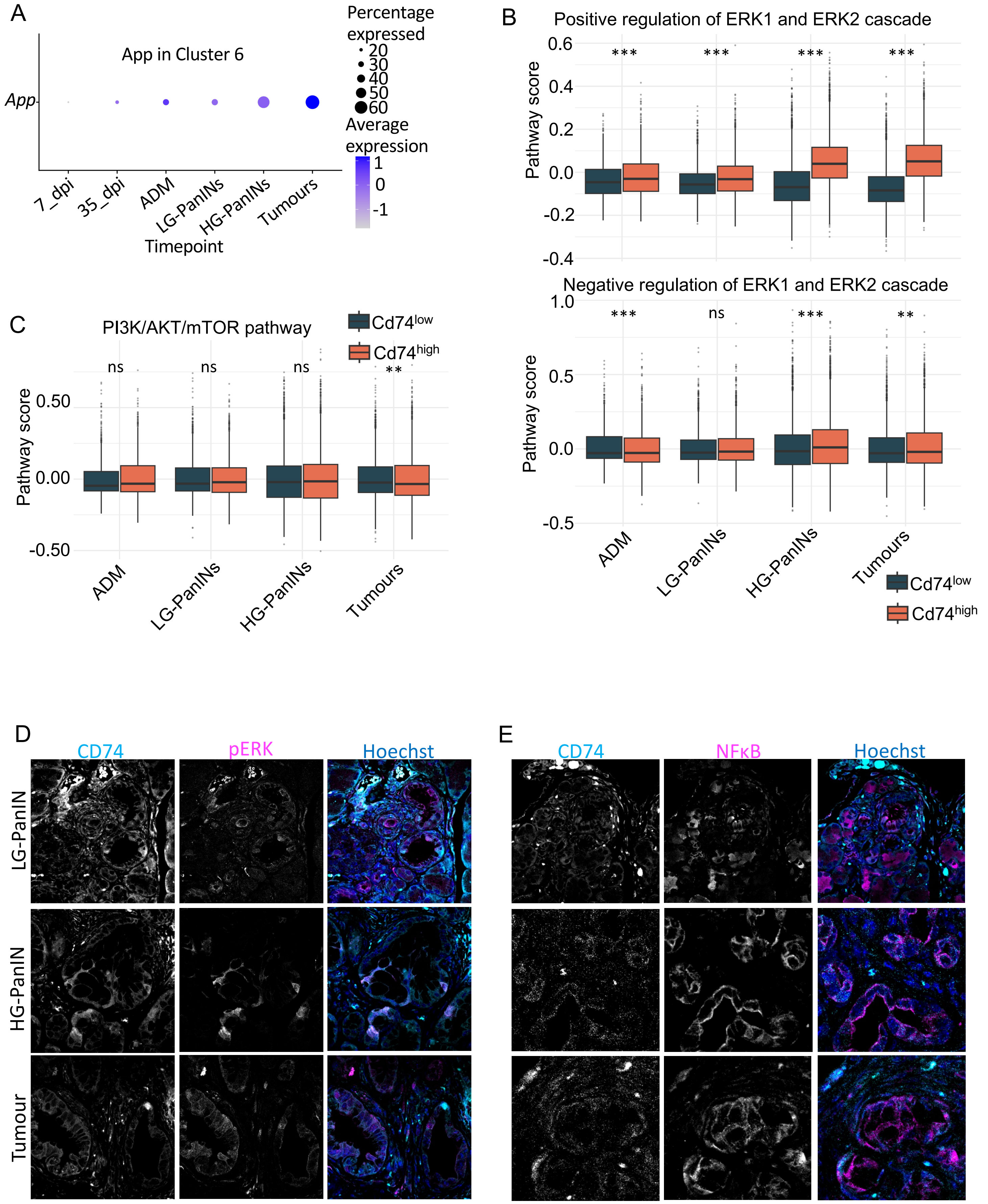
A
App in Cluster 6
Percentage expressed
20
30
40
50
60
Average expression
1
0
-1
App
7_dpi
35_dpi
ADM
LG-PanINs
HG-PanINs
Tumours
Timepoint
B
Positive regulation of ERK1 and ERK2 cascade
Pathway score
0.6
0.4
0.2
0.0
-0.2
-0.4
***
***
***
***
Negative regulation of ERK1 and ERK2 cascade
1.0
0.5
0.0
-0.5
***
ns
***
**
ADM
LG-PanINs
HG-PanINs
Tumours
Cd74low
Cd74high
C
PI3K/AKT/mTOR pathway
Cd74low
Cd74high
ns
ns
ns
**
Pathway score
0.50
0.00
-0.50
ADM
LG-PanINs
HG-PanINs
Tumours
D
CD74
pERK
Hoechst
LG-PanIN
HG-PanIN
Tumour
E
CD74
NFκB
Hoechst

**Sup Figure 9**

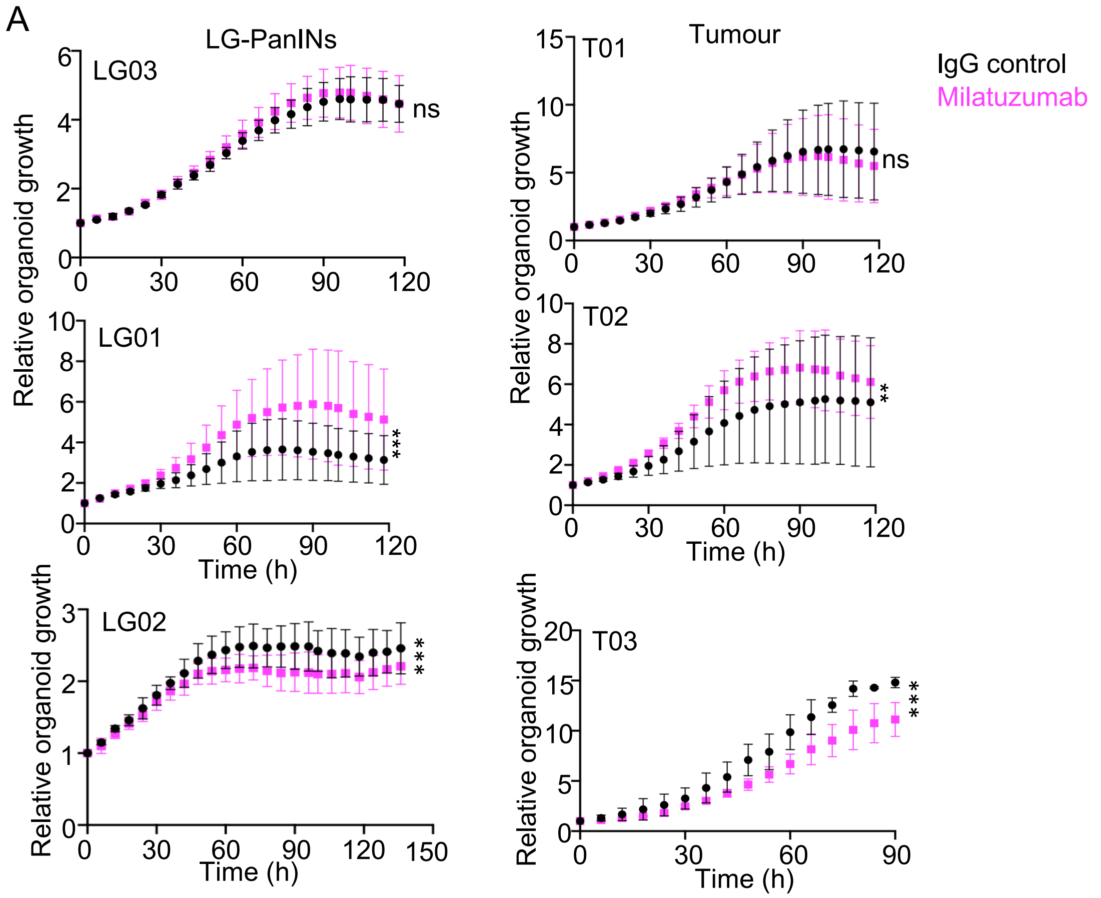
A
LG-PanINs
Tumour
IgG control
Milatuzumab
LG03
ns
LG01
***
LG02
***
T01
ns
T02
**
T03
***
Relative organoid growth
Time (h)

**Figure S1. Defining experimental timepoints.**

(A) Representative images of alcian blue staining (top), Mucin 1 and Mucin 5AC staining and endogenous RFP fluorescence. Scale bar: 100µm. Dashed lines show ADM examples. White arrow heads point at Mucin 1 positive cells. (B) Quantification of alcian blue positive lesions per tissue area at 8, 11, 14 and 17 weeks post tamoxifen induction (n= 3 mice/timepoint). (C) Percentage of LG-PanINs (Mucin 5AC positive) or HG-PanINs (Mucin 5AC and Mucin 1 positive) at each timepoint (n= 3 mice/timepoint). (D) Percentage of RFP fluorescence per total tissue area. N=mouse numbers described in the graph with average values/mouse represented by a data point. Data represent mean ± SD.

**Figure S2. Single cell/nuclei transcriptomic data annotation.**

(A-B) UMAP visualizations of pancreatic scRNAseq data including all cell types and timepoints (Figure1C) coloured according to log-normalized gene expression level for each cell. (A) Plots show Neuroendocrine marker *Resp18*, Pericyte marker *Rgs5*, Immune cell markers *Ptrcp*, *Cd19* (B cells), *Cd4* (T cells), *Cd68* (macrophages), Fibroblast marker *Col1a1*, Endothelial marker *Pecam1*. (B) UMAP plots show pancreatic epithelial cell markers including Acinar marker *Cpa1*, Ductal marker *Krt19*, Metaplastic markers *Krt19* and *Clu* and Neoplastic markers *Arg2*, *Muc5ac* and *Tff1*. (C) UMAP plots containing exocrine epithelial cells (Figure 1D) and all timepoints coloured according to log-normalized gene expression level for each cell. Plots show Acinar markers *Cpa1* and *Cpa2*, Ductal marker *Krt19*, Metaplastic markers *Krt19*, *Sox9* and *Clu*, Neoplastic markers *Arg2*, *Muc5ac* and *Tff1*, Gastric-like markers *Muc5ac*, *Gkn1* and *Gkn2*, Tuft markers *Dclk1* and *Pou2f3* and Progenitor marker *Msn*.

**Figure S3. Clusters 5 and 6 express higher progenitor signatures than other exocrine clusters.**

(A) Bubble plot of pancreatic stem/progenitor marker genes across epithelial exocrine cell types at all timepoints showing the percent of cells in each cluster expressing the gene and the log-averaged expression value. The plot shows the percent of cells in each cluster expressing the gene (bubble size) and the log-averaged expression value (colour scale). (B) Waterfall plot showing mean score for Pancreatic Cancer Stem Cell signature (PCSC; Figure 1F) comparing exocrine clusters. Wilcoxon tests were used to analyse the data. ***P < 0.0001. (C) Representative confocal images of mouse tissue at different timepoints stained with stem/progenitor markers in A, including Cytokeratin 19 and CD44. Scale bar: 20µm. (D) Bar plot showing the percentage of cells per exocrine cluster in each timepoint. (E) Gene set scores of cluster 6 (brown plots) compared to all other exocrine clusters (blue plots) for Malta Curated Stemness Markers (left) and Ramhlo Stemness UP in all timepoints.

**Figure S4. Gene signatures previously described as pancreatic progenitor-like state and malignant programme align with Clusters 5 and 6.**

(A-B) Bubble plots of (A) progenitor-like cell state[13] and (B) MP10[21] marker genes across epithelial exocrine cell types at all timepoints showing the percent of cells in each cluster expressing the gene and the log-averaged expression value. The plot shows the percent of cells in each cluster expressing the gene (bubble size) and the log-averaged expression value (colour scale).

**Figure S5. EPCs (cluster 6) overexpress genes involved in stress response, inflammatory response, cell survival and cell-cell and cell-ECM attachment.**

(A) Bubble plots of top Differentially Expressed Genes (DEGs) in cluster 6 versus all other exocrine clusters over time showing the percent of cells in each cluster expressing the gene (bubble size) and the log-averaged expression value (colour scale). (B, C) Scatter plot showing percentage of total Integrin β6 (B) and Claudin 4 (C) highly expressing cells in LG-PanINs, HG-PanINs and tumours. Each dot represents an individual mouse, and the graph shows the median. (D-F) Dot plots showing significantly enriched Reactome pathways (D), Wikipathways (E) and Gene Ontology Biological Processes (F) in cluster 6 compared with other exocrine clusters. Gene Ratio shows the proportion of genes associated with each pathway. Dot size indicates the number of genes contributing to each pathway (Count), while dot colour represents the adjusted p-value (Benjamini–Hochberg correction). Only top 20 significant pathways with an adjusted p-value < 0.05 are shown. RTK: receptor tyrosine kinase; IGFBPs: insulin-like growth factor binding proteins; GMS: glomerulosclerosis; Reg: regulation.

**Figure S6. Cluster 6 (EPCs) shows progenitor potential in pseudotime analysis.**

(A) Violin plot showing general pseudotime values at each timepoint including all exocrine epithelial cells together. (B) Box plot showing pseudotime values per clusters including all timepoints in each cluster. (C) Heatmap showing the activity of gene modules identified using Monocle3 find_gene_modules() analysis across the exocrine cell clusters. Colour indicates the scaled module activity score. Hierarchical clustering groups modules and clusters according to similarities in their activity profiles. (D) Dot plot showing significantly enriched Gene Ontology Biological Processes in Module 1 compared with other modules. Gene Ratio shows the proportion of genes associated with each pathway. Dot size indicates the number of genes contributing to each pathway (Count), while dot colour represents the adjusted p-value (Benjamini–Hochberg correction). Only pathways with an adjusted p-value < 0.00001 are shown.

**Figure S7. Few PanINs/GLS are enriched for Sca-1 positive cells.**

(A) Representative confocal images of mouse tissue stained with Sca-1/Ly6a antibody at different timepoints. Scale bar: 50μm. (B) Scatter plot showing percentage of total Sca-1/Ly6a expressing cells in LG-PanINs, HG-PanINs and tumours. Each dot represents an individual mouse, and the graph shows the median. (C) Bar plot showing the percentage of PanINs/GLS containing different proportions of epithelial cells showing positive staining for Sca-1. Data represent mean ± SD. N=3 mice per timepoint and a 5-12 PanINs/GLS per mouse.

**Figure S8. Only few PanINs/tumours show a high proportion of cells expressing stress response markers.**

(A-C) Scatter plot showing percentage of total Nupr1 (A), Clusterin (B) and phospho-EphA2 (pEphA2; C) highly expressing cells in LG-PanINs and HG-PanINs. Each dot represents an individual mouse, and the graph shows the median. (D) Bar plots showing the percentage of PanINs/GLS containing different proportions of epithelial cells showing positive staining for pEphA2 (E). Data represent mean ± SD. N=3 mice per timepoint and a minimum of 10 PanINs/GLS per mouse. (E) Representative confocal images of mouse tissue stained with pEphA2 and E-cadherin at different timepoints. Scale bar: 50μm. (F) Representative confocal images of Nupr1 staining in HG-PanIN organoids.

**Figure S9. Cd74 expressing exocrine cells overexpress readouts of ERK1/ERK2 and NFκB pathway.**

(A) Bubble plot of *App* expression in cluster 6 along disease progression. Dot size represents the percent of cells in each cluster expressing the gene and the colour the log-averaged expression value. (B, C) Box plots showing pathway scores for the GOBP_positive_regulation_of_ERK1_and_ERK2_cascade (B top), GOBP_negative_regulation_of_ERK1_and_ERK2_cascade (B bottom) and Hallmark_Pi3k_AKT_mTOR_fignalling (C) in $Cd74^{high}$ (top quartile; orange) and $Cd74^{low}$ (bottom quartile; blue) exocrine cells at each timepoint. Wilcoxon tests were used to analyse the data. ns=not significant; P > 0.05; **P < 0.001; *** P > 0.0001. (D, E) Representative confocal images of mouse tissue stained with CD74 and pERK (D) or CD74 and NFκB (E) antibodies at different timepoints. Scale bar: 20 μm.

**Figure S10. Organoid response to CD74 blockade is variable in LG-PanIN- and tumour-derived lines.**

(A) Graphs showing organoids growth over time in three independent LG-PanIN-and three independent Tumour-derived organoid lines (3 independent experiments per line) treated with IgG1 or Milatuzumab 5μg/ml for 20-148h. Data represent mean ± SD. ANOVA tests were used to analyse the data. ns=not significant; P > 0.05; **P < 0.001; *** P > 0.0001.
